# Effects of fire regime on the bird community in an Amazonian savanna

**DOI:** 10.1101/2022.08.01.502386

**Authors:** Laís Araújo Coelho, Camila Duarte Ritter, Albertina Pimentel Lima, Renato Cintra, William E. Magnusson, Tânia Margarete Sanaiotti

**Affiliations:** Department of Ecology, Evolution, and Environmental Biology, Columbia University, MC-5557, 1200, Amsterdam Avenue, New York, NY 10027, USA; Instituto Nacional de Pesquisas da Amazônia. Avenida Efigênio Sales, 2239, Manaus, AM 69060-001, Brazil; Instituto Juruá, Rua das Papoulas, 97, Aleixo, 69083-300, Manaus, AM, Brazil

**Keywords:** Alter do Chão, bird community, fire extent, fire frequency, forest-savanna boundary, Tropical America

## Abstract

Savanna ecosystems are maintained by fires with a fire-adapted biota, and savannas occur in Amazonia in patches surrounded by tropical forest. Different fire regimes can generate structurally diverse vegetation, and the composition of savanna bird assemblages is known to be closely related to vegetation structure. However, long-term approaches and interaction of fire with other environmental factors need to be explored for the better understanding of the effects of fire on birds. In an Amazonian landscape composed by savanna and forest, we investigated the effects of different fire regimes in a 12-ha area in three periods through 23 years. We also examined the effects of frequency and extent of fires, tree cover, and distance to forest on bird composition in twelve 3.7- ha savanna plots. Birds were surveyed with mist-nets and species were classified as to their habitat use by comparison of registers in forest and savanna plots through visual/acoustical surveys. After 13 years without fire, many forest species colonized the area and some savanna species were lost. Fire regime affected avifauna assemblages. The avifauna was sensitive to the occurrence of fires. After one fire event in a plot that had not burned for 12 years, some savanna species returned. These results highlight the effects of the fire regime on birds and indicate that many savanna bird species depend on the occurrence of regular fires.

## Background

Savanna vegetation is characterized by the occurrence of grasses, shrubs, and trees with discontinuous canopy cover (Ferreira et al. 2015, Silva et al. 2019; Azevedo et al. 2020). Savannas differ from other open vegetation types in that they burn regularly (Ramos-Neto and Pivello 2000). In Tropical America, there are three main domains of the savanna biome: the Llanos and the Guianan domains in northern South America, and the Cerrado domain in central South America (da Silva and Bates 2002, Furley 2006, Azevedo et al. 2020). Beyond these domains, there are also several smaller areas of savanna scattered across Tropical America, such as in the Atlantic Forest, in Central America, on Caribbean islands, and within lowland Amazonia (de Carvalho and Mustin 2017, Pennington et al. 2018). In the Amazon, savannas are estimated to cover between 1.8% (de Carvalho and Mustin 2017, Mustin et al. 2017) and 4% (Pires and Prance 1985) of the region and occur isolated in patches of different sizes, surrounded by a matrix of continuous canopy and moist dependent tropical rainforest (Huber 1982, de Carvalho and Mustin 2017).

Savannas of tropical humid regions are maintained mainly by the incidence of fires (Moreira 2000, Bond and Keeley 2005). Fire regimes differ in the combination of several factors, such as frequency, intensity, extent, and the period of year in which they occur (Bond and Keeley 2005), and some plant species show adaptations to this disturbance (Fidelis and Zirondi 2021). Savanna organisms are resilient to milder fires, similar to natural fire regimes, and are impacted by fires that are more frequent or that occur in seasons different from the natural fire period (Griffiths and Christian 1996, Ramos-Neto and Pivello 2000, Medeiros and Miranda 2008, Smit et al. 2010, Fidelis and Zirondi 2021). Savannas are threatened by changes to the natural fire regimes; while in some areas fires have become more frequent, other savannas undergo a process of densification of woody vegetation, possibly due to fire suppression (Sanaiotti and Magnusson 1995, Medeiros and Miranda 2008, Moustakas et al. 2010, Lima et al. 2020).

The effects of the alteration in natural fire regimes could threaten the Amazonian savanna biota. However, studies on the effect of fires on the savanna biota are generally incomplete, fragmented, and frequently based on controlled experiments that do not reflect the real variation of the effects of non-experimental fire regimes (Parr and Chown 2003, Russell-Smith et al. 2010). Amazonian savannas have a unique biota and also allow the expansion of the Cerrado biota into the region, including some endangered species (Aleixo and Poletto 2007), connecting populations of Llanos and the Guiana domains with the Cerrado (Ritter et al. 2021), and maintain genetic diversity because they have more recent diversification times than populations of the main domains (van Els et al. 2021). Despite their limited distribution, importance for biota, and pressure from the rapid expansion of agriculture and livestock, the Amazonian savannas are underrepresented in protected areas (Sanaiotti and Magnusson 1995, Santos and Silva 2007). In this context, studies on well-known taxonomic groups can offer critical insights for the savanna dynamics under different levels of disturbance.

Birds have well documented geographic and morphological variability. The composition of the savanna avifauna is associated with the severity of the fires and varies along the gradient of disturbance (Barlow and Peres 2004, Clavero et al. 2011, Davis and Miller 2018). However, evidence of diversified responses to fire regimes led to “pyrodiverse” management techniques in savannas, without regard to the ecological significance of some fire patterns (Parr and Andersen 2006). The investigation of the responses of birds to different intensities and frequency of fires in areas of contact between savanna and forest is important in order to understand the effects of forest expansion or of the use of fire for the restoration of savanna avifauna (Artman et al. 2005, Moura et al. 2016, Davis and Miller 2018). In addition, long-term approaches and those that consider the spatial variation of fires and their interaction with other environmental factors need to be incorporated for a better understanding of the relationship between fire and fauna (Parr and Andersen 2006, Driscoll et al. 2010).

We analysed variation in bird assemblages in an eastern Amazonian savanna in response to different spatial and temporal scales of fires events. We used two distinct approaches to evaluate the effects of fire, one based on a time sequence of changes within plots, and another based on spatial differences in fire events history. We evaluated the dynamics of the avifauna over 23 years, in which 10 were regularly burned and the last 13 years without fire. Due to the limited sampling periods for the temporal analysis and as the area had burned before the study began, it was not possible to include a strict control/impact design. We then, used 12 widely dispersed plots to evaluate the effects of the frequency and extent of fires, tree cover, and distance to forest on bird assemblages, to complement time variation in bird assemblages with spatial variation that included plots with distinct fire histories (Legendre and McArdle 1997). We hypothesised that (i) avifauna composition would be significantly different after 13 years without fire, with colonization by some forest species; (ii) fire frequency and extent would affect avifauna composition, with a higher abundance of savanna species in plots with higher occurrence and extent of fires.

## Material and Methods

### Study area

The study was conducted in an enclave of savanna within forest in the state of Pará, Brazil, in the municipality of Santarém, near the village of Alter do Chão (2 ° 31 ’S, 55 ° 00’ W) on the right bank of the Tapajós River (Figure 1). The area covers about 10,000 ha distributed in four non-contiguous enclaves, with a larger portion of the area being composed of sandy soil covered by an herbaceous layer of varying height and density, a shrub layer of 60-80 cm, and a tree layer up to 10 m (Cintra and Sanaiotti 2005, Magnusson et al. 2008). The savanna contains different-size patches of semi-deciduous forest and is surrounded by the same vegetation (Miranda 1993, Cintra and Sanaiotti 2005). The climate has two well-defined seasons: a dry season from July to December, and a rainy season from January to June, with 75% of rainfall occurring from December to June. Fires are frequent in the area; local people traditionally burned the savannas regularly, usually during the dry season, but environmental agencies have discouraged burning during the past few decades, suppressing fires in many areas.

**Figure 1.**
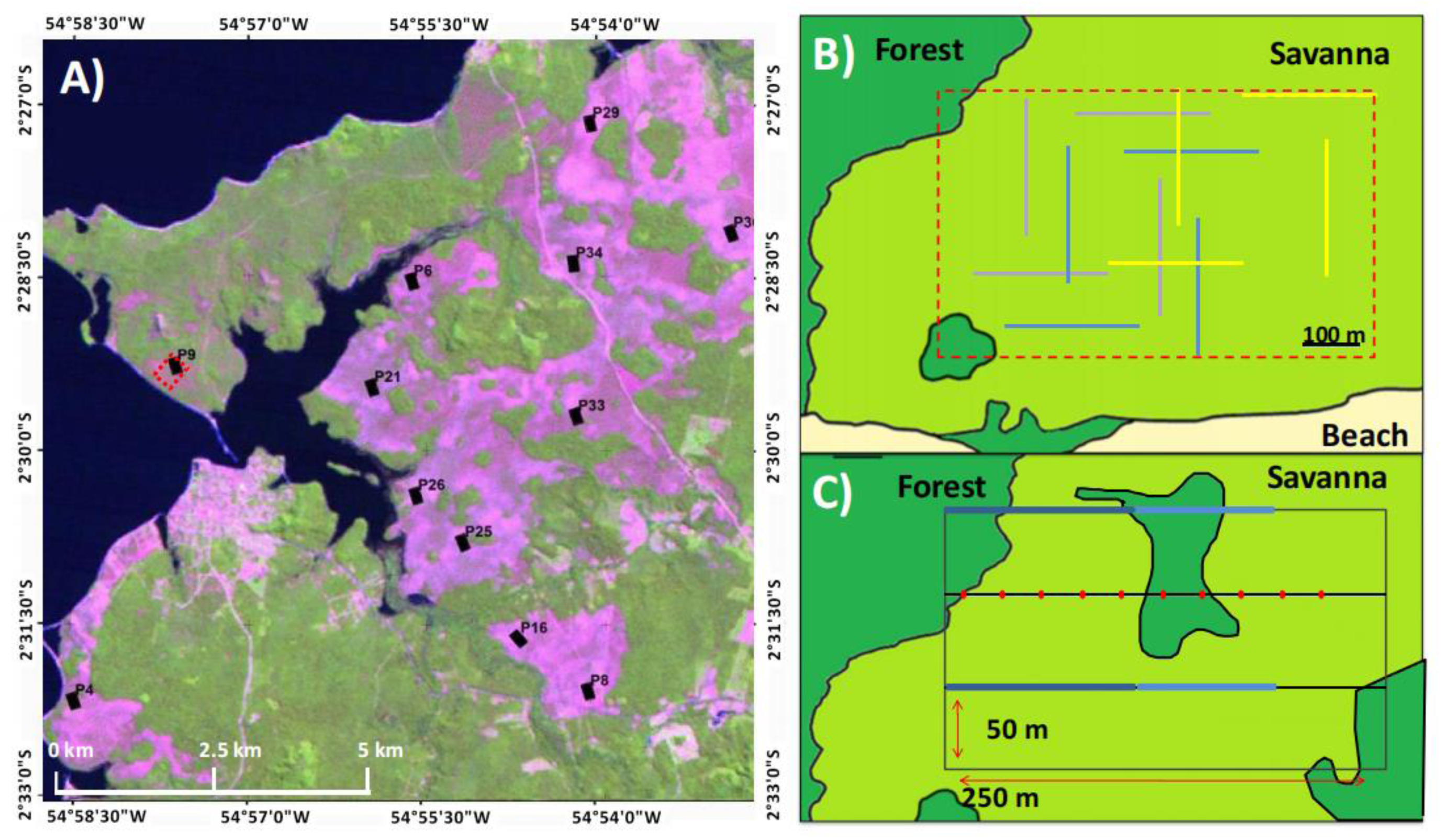
A) Landsat image of the study area from June 2010. The purple portion represents the savanna area; the green corresponds to forest vegetation; and the black colour is the Tapajos River. The black rectangles represent the 12 sampling plots (150 x 250m), while the single red-dotted rectangle indicates the area where the long-term avifauna dynamics study was undertaken (12ha, “peninsula”); B) Sampling design of the temporal analysis of variation in bird community study. The outer red dashed rectangle represents the sampling site. The solid lines represent net-lines, and each set of 4 lines in the same color represents net-line positions in one sample period (in this case, the sampling schemes of three samples are represented). The overlap of the positions of net-lines throughout the study formed a grid of 12 ha (outer red dashed rectangle); C) Sampling design of the spatial analysis of bird assemblages. Four parallel transects of 250 m, distant 50 m from each other, were placed in each plot. In the all four transects, for each 2 m the fire extension and vegetation cover percentage were measured (sum of points with burnt vegetation or wood vegetation or canopy coverage / total points), represented by the red dots. Birds were sampled in two net-lines placed in the first and third transects represented by blue lines (dark blue represents the six mist-nets of 12 m and light blue the nine mist-nets of 12 m).

### Bird sampling

We captured birds with mist-nets (mesh size 36 mm). We identified the birds based on identification guides (Mata et al. 2006, Ridgely and Tudor 2009) and consultation with specialists. The nomenclature of birds follows the “List of Brazilian Birds” of the Brazilian Committee of Ornithological Records (CBRO 2011). Birds captured in this study were marked with metal bands provided by CEMAVE-ICMBio (authorization number 1177 / 4, ICMBIO / SNA) or coloured plastic bands.

### Temporal analysis

We captured birds in three sampling periods over 23 years (1987/88; 1996/97 and 2009/10): five samples in the period from September 1987 through August 1988, three samples in the period from September 1996 through February 1997 and five samples in the period from August 2009 through June 2010. The sampling site was a savanna area of about 12 ha (250 x 450 m) located on a peninsula (Figure 1A, dotted red rectangle). From at least 1985 until 1997 the area was under a regular fire regime (Sanaiotti and Cintra 2001). From 1998 until 2010 there were no fires in the study site. The sampling protocol was standardized throughout the whole study period, in which we used 20 mist nets of 12 x 2.5 m (36 mm mesh size). The nets were positioned consecutively, forming a single straight mist-net line. Every second day, the location of the mist-net line was changed to a perpendicular position distant up to 100 m from the previous net line (Figure 1B). We varied the net-line positions between samples within periods, to cover the maximum possible area of the 12-ha site (Fig. 1B). Nets were opened from 06:00 to 09:00 and from 15:30 to 17:30. Each sample consisted of four positions of net lines, in which we considered the four morning samples of each net-line position and the two first sampled afternoon periods, accruing a total of 320 net-hours per sample. The captures and recaptures are shown in Table S1.

### Spatial analysis

To assess the spatial variation in bird assemblages, we sampled twelve 3.75 ha plots that were separated by at least 1 km. Each plot had four parallel 250 m transects spaced 50 m apart (Fig. 1C). For bird sampling, we used two of the transects spaced 100 m apart, each with 15 mist nets (36 mm mesh size, nine 9 x 2.5 m mist nets and six 12 x 2.5 m mist nets), in a total of 30 mist nets per plot. The mist nets were opened in the same time interval described previously for one day in each plot. Sampling was done in the dry season from August to October 2010.

### Additional short-term fire-effect observations

Plot 9, which had not burned since 1997, had an unexpected fire on 23 October 2010. This plot was sampled again in November 2010, immediately following the fire event, with the same protocol used in the 12 plots.

### Bird classification

We classified bird species regarding the habitat in which they were most common in the savanna-forest mosaic of the study area. In 1999 and 2000, RC undertook visual and acoustic sampling of birds in 22 savanna plots and 22 forest plots (distributed in isolated forest fragments in savanna matrix and in the adjacent continuous forest) for a posterior classification of species as to their habitat use. The 22 savanna plots had the same dimensions as the 12 plots sampled with mist-nets in this study. RC sampled two 250 m transects from 6 to 9 am and from 4 to 6 pm in the afternoon, stopping for two minutes every 50 m and registering every bird heard or seen within a 10 m radius (Cintra and Sanaiotti, 2005; Cintra, Magnusson and Albernaz, 2013). The abundances of the species in the 44 savanna and forest plots were compared with a Wilcoxon test. The species that were more abundant in forest plots (p<0.05) were classified as forest species (F), those that were more abundant in savannas were classified as savanna species (S), and species whose abundances did not differ significantly between the two habitats were classified as indifferent (I). Six species (Table 1) were classified based on the literature because they were not recorded in point-count surveys in any of the 44 plots.

**Table 1.** Species captured by mist-net sampling in the savanna of Alter do Chão–-PA. Species marked with • had not been registered before in the study area (Sanaiotti & Cintra, 2001). Habitat use refers to the habitat where the species was most frequent in the region (Savanna, Forest or Indifferent), * are species classified based on literature, for which we did not have enough data for analyses.

| Family | Species | Habitat Use |
| --- | --- | --- |
| **Columbidae** | *Crypturellus parvirostris* | S |
|  | *Columbina passerina* | S |
|  | *Claravis pretiosa* | I |
|  | *Zenaida auriculata* | S* |
|  | *Leptotila rufaxilla* | F* |
| **Bucconidae** | *Nystalus maculatus* | S |
| **Dendrocolaptidae** | *Dendroplex picus* | F |
|  | *Lepidocolaptes angustirostris* | S |
| **Thamnophilidae** | *Formicivora grisea* | F |
|  | *Formicivora rufa* | S |
|  | *Thamnophilus stictocephalus* | F |
| **Pipridae** | *Neopelma pallescens* | F |
|  | *Manacus manacus* | F |
| **Tityridae** | *Pachyramphus polychopterus* | F |
|  | *Pachyramphus rufus*• | F* |
| **Tyrannidae** | *Tolmomyias flaviventris* | F |
|  | *Todirostrum cinereum* | F |
|  | *Camptostoma obsoletus* | S* |
|  | *Hemitriccus striaticollis* | F |
|  | *Elaenia flavogaster* | S |
|  | *Elaenia cristata* | S |
|  | *Elaenia chiriquensis* | S |
|  | *Elaenia parvirostris* | S* |
|  | *Phaeomyias murina* | S* |
|  | *Suiriri suiri* | S |
|  | *Myiopagis viridicata*• | F* |
|  | *Myiarchus swainsoni* | I |
|  | *Myiarchus ferox* | S |
|  | *Myiarchus tyrannulus* | S |
|  | *Sirystes sibilator* | F |
|  | *Megarynchus pitangua* | F |
|  | *Tyrannus albogularis* | I |
|  | *Tyrannus melancholicus* | I |
|  | *Griseotyrannus aurantioatrocristatus*• | S* |
|  | *Empidonomus varius* | I |
| **Vireonidae** | *Cyclarhis gujanensis* | S |
|  | *Vireo olivaceus* | F |
|  | *Hylophilus semicinereus*• | F* |
|  | *Hylophilus pectoralis* | F |
| **Hirundinidae** | *Stelgidopteryx ruficollis* | I |
| **Troglodytidae** | *Troglodytes musculus* | S |
|  | *Cantorchilus leucotis* | F |
| **Turdidae** | *Turdus leucomelas* | F |
| **Thraupidae** | *Saltator maximus* | I |
|  | *Tachyphonus rufus* | F |
|  | *Ramphocelus carbo* | F |
|  | *Tangara cayana* | S |
|  | *Thraupis episcopus* | I |
|  | *Thraupis palmarum* | F |
|  | *Schistochlamys melanopis* | I |
|  | *Cyanerpes cyaneus* | F |
| **Emberizidae** | *Ammodramus humeralis* | S |
|  | *Volatinia jacarina* | S |
| **Cardinalidae** | *Piranga flava* | S |
| **Fringilidae** | *Euphonia chlorotica* | I |

### Environmental variables

The fire-history data from each plot was collected from 1997 to 2009, except for 1998. The four 250 m transects in the plots were sampled annually after the burning period (usually from August to December). To estimate the burn extent in a plot, we checked for burnt herbaceous vegetation (which regenerates within months after fire) at 2 m intervals, and the annual burn extent value was the percentage of burnt points per plot (number of points with burnt vegetation/total plot points). Fires were spatially congruent, so there were no points on burned ground with unburned vegetation around. The fire history of the plots was calculated by summing all of the annual burn extent for each plot (range: 0 – 1,200). Another fire-history variable that we considered was the frequency of fires regardless of their extent. This variable was calculated by summing the number of years throughout the 12 sampling years in which the plot was burned. Therefore, this value ranged from 0 to 12 and will hereafter be referred to as fire frequency.

Data on tree cover were collected from July to August 2007. We used tree cover measured 3 years prior (with no fire occurrence) to bird sampling because tree cover did not change significantly in a four year interval in the study region (Vasconcelos et al. 2008), or during subsequent years (personal observation). At 2-m intervals along the four transects in each plot, a cylindrical metal stick 1-m long with 2 mm diameter was positioned perpendicularly to the ground and it was noted whether it touched a tree or was under a tree canopy (height > 2m). We considered tree cover as the proportion of points that met this criterion. For more information about the method, see Magnusson et al. (2008).

The composition of the bands 1, 2, 3, 4, 5, and 7 of a Landsat TM 5 image (orbit 227, point 62) of 29 June 2010 was georeferenced to a geographic information system (ESRI ® ArcMap ™ 3.9) using paths and points collected with GPS in the study area. The resulting image composition underwent a process of supervised classification by maximum likelihood with the program ENVI 4.5 © 2008. This resulted in a thematic map of the study area containing the following classes of land cover: 1) savanna, 2) forest, 3) water bodies, and 4) other. We obtained the distance to forest for each plot from this composition, considered as the shortest distance measured from the centroid of each plot and the edge of the closest forest.

### Data analyses

We did not include nightjars or hummingbirds in the analyses because our mist-net sampling was not appropriate for capturing these groups. Four species (*Myiarchus swainsoni, Myiarchus ferox, Phaeomyias murina,* and *Camptostoma obsoletum*) were not included in the analysis of temporal variation because irregularities in their identification were detected. These species were included in the analysis of spatial variation given that, during the sampling of this data, birds were photographed for posterior identification. Recaptures within each temporal or spatial sample were not included in the analysis. We used the number of captures of each species as an index of relative abundance for each sampling unit for all analysis. All statistical analyses were done in R software (R Core Team 2021).

To test the first hypothesis (avifauna composition would be significantly different after 13 years without fire, with colonization by some forest species), we used a two-dimensional non-metric multidimensional scaling (NMDS) ordination as implemented in the metaMDS function in the R package vegan 2.5-7 (Oksanen et al. 2010), with the data from the peninsula over the 23 years. Compositional dissimilarity was quantified with the Bray-Curtis dissimilarity index. We tested the significance of the NMDS axis for the year of sampling through the envfit function. Next, we used the Betadisper analysis followed by the TurkeyHSD test to check for difference among group variances. As no difference was found (p > 0.84) for any pairwise comparison, we used Permutational Analysis of Variance (PERMANOVA) to assess whether sampling period affected bird assemblage composition. To test which sampling periods were significantly different from others, we used the function “pairwise_adonis” with the package ranacapa v. 0.1.0 (Kandlikar 2021)(Kandlikar 2021) with Bonferroni correction. To test if forest species were more abundant in the no-fire period and savanna species more abundant in the fire period, we used a Kruskal-Wallis test for both groups separately followed by a pairwise Wilcox test to detect significantly different groups. To visualize the change in bird community composition, we used a direct ordination of samples (McCune et al. 2002) in chronological order to view changes in the community throughout the sampling period in a compound graph in which the relative abundance of all species is shown (Dambros 2014)(Dambros 2014).

To test the second hypothesis (fire frequency and extent would affect avifauna composition, with a higher abundance of savanna species in plots with higher occurrence and extent of fires), we used the relative-abundance dissimilarity measured with the Bray-Curtis distance and the environmental variables (except for plot 9 after fire since the variables were collected before the fire event) that were standardized to mean = 0 and variance = 1. Most variables had high Pearson correlations, except for forest distance (Table S2). Because fire frequency, fire extent, and tree coverage were correlated, we used the community dissimilarity as the response variable and the fire frequency and the distance to forest as explanatory variables in a PERMANOVA. PERMANOVA analyses of fire extent and tree coverage are presented in the Supplementary html file.. Additionally, we tested the effects of environmental variables on overall bird abundance and species richness of savanna species and forest species separately. We used generalized linear models (Crawley 2007) with Poisson error distribution after checking for the distribution of residuals. We defined a set of models to explain variation in abundance and richness for savanna species and for forest species separately using fire frequency and distance to the forest separately and together in the same model. The final model set included an additional constant, intercept-only model, comprising a total of four models for each dependent variable. Models were selected using an information-theory approach based on AIC (Akaike 1974) using the corrected AIC (dAICc) for small with the bbmle v. 1.0.25 package (Bolker and Bolker 2017). Models including fire extent and tree coverage separately and with distance to the forest as an additive effect are presented in the supplementary html material.

For the additional short-term fire-effect observations, to visualize the change in the bird composition in plot 9, we plotted the inverse of abundance pre-fire and total abundance post-fire of each species in a lollipop graph with ggplo2 v. 3.3.5 (Wickham 2016). Additional R packages used for data curation and visualization were tidyverse v.1.3.1 (Wickham 2017), and plotly v.4.10.0 (Sievert 2020). The html with the tables, code, and results are available as supplementary material.

## Results

### Temporal analysis

A total of 620 individuals of 46 species were caught in 723 captures in 13 sampling occasion (five in 1987/88, three in 1996/97, and five in 2009/10) on the peninsula. Our first hypothesis, that avifauna composition would be significantly different after 13 years without fire, with colonization by some forest species, was supported. The samples collected after 13 years without fire did not overlap with the samples collected under the frequent-fire regime in the space defined by the two NMDS axes (Figure 2, stress = 0.13; envfit: r^2^ = 0.59, p = 0.002). The first NMDS axis split samples from the 2009-10 period without fire (green circles) from the two sampling periods under fire regime (1987-88 [red triangles] and 1996-97 [orange squares]), which overlapped along the NMDS axis (Figure 2). The results of PERMANOVA also indicated that the bird assemblage composition varied significantly among the three sampling periods in the 12-ha study site of the peninsula (*F* = 2.73, *P* = 0.002). The pairwise comparisons showed that 1987-88 and 1996-97 bird assemblages were not significantly different, but both were significantly different from the 2009-10 bird assemblage (Table 2). Forest species were more abundant in the period without fire (Chi-square = 17.54, p < 0.001), but did not differ between 1987/88 and 1996/97; both differed from 2009-10 (Table S3). However, savanna species were not significantly different among sampling periods.

**Figure 2.**
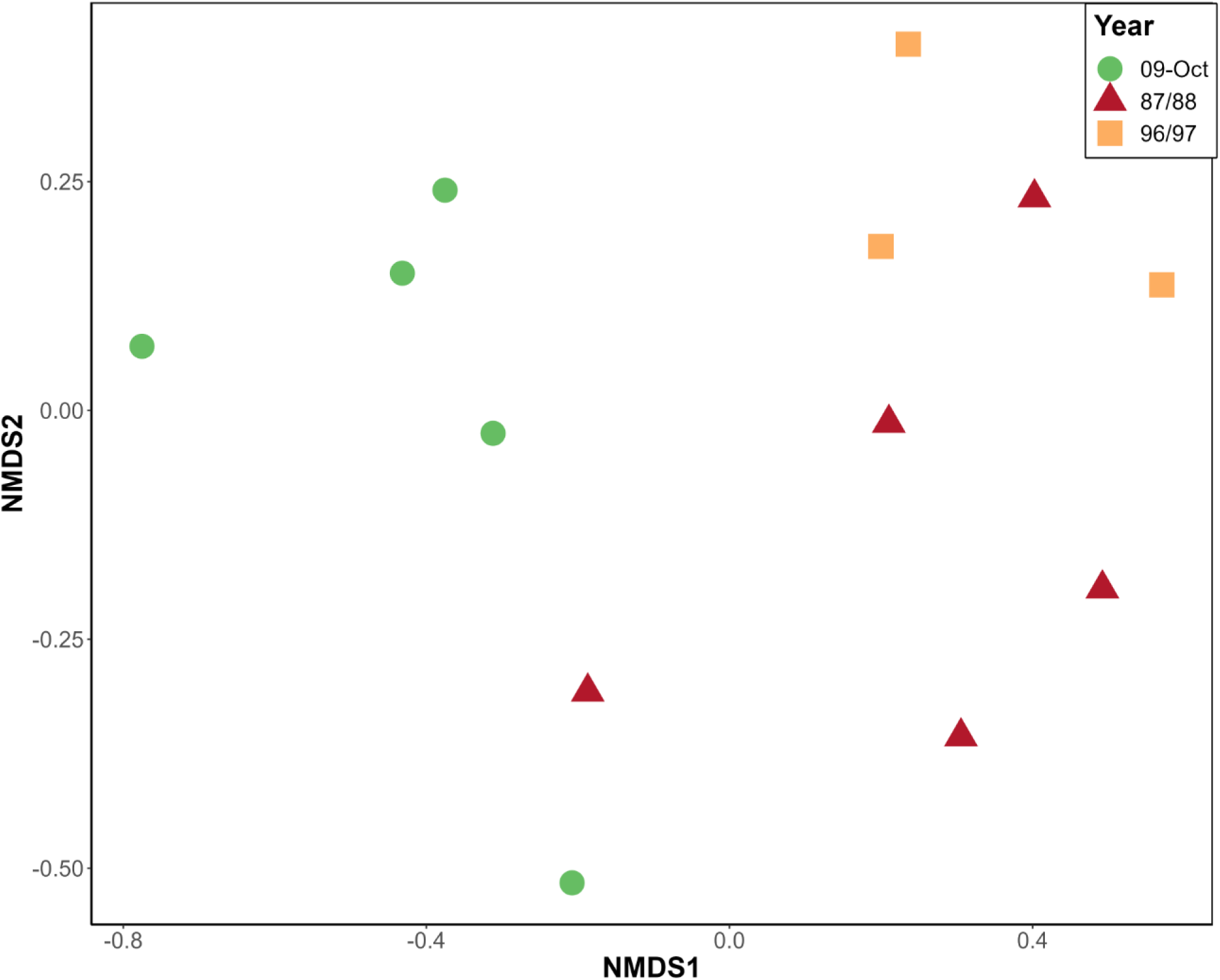
Non-metric Multidimensional Scaling (NMDS) with long-term samples (stress = 0.13). The red triangles and orange squares represent samples from periods when the 12-ha study site on the peninsula was under a regular fire regime. The green circles represent samples collected after 12 years without fire. The first NMDS axis split without overlap the samples under regular fire regime period from the samples in the period without fire.

**Table 2.** Pairwise Adonis results from the temporal Permutational Analysis of Variance – PERMANOVA (*F* = 2.73, *P* = 0.003). In bold are hig<u>hlighted the pairwise significantly different bird assemblage after B</u>onferroni correction.

| **Pairs** | **F.Model** | **R2** | **p.value** | **p.adjusted** |
| --- | --- | --- | --- | --- |
| 1987/88 vs 1996/97 | 1.265589 | 0.174189 | 0.212 | 0.64 |
| **1987/88 vs 2009/10** | **3.45626** | **0.301692** | **0.009** | **0.03** |
| **1996/97 vs 2009/10** | **3.232536** | **0.350124** | **0.017** | **0.05** |

### Spatial analysis

In the 12 plots sampled in August 2010, excluding the resampling of plot 9, we caught 666 individuals belonging to 44 species in 744 captures. There were no recaptures between different plots. Plot 9 was located in an area that had not burned for 12 years (the last fire was in 1997), while the fire frequency among the other plots ranged from 6 to 10 fires in the last 12 years (Table 3). Ten plots (except for 4 and 9) were burned in 2009, one year before the bird sampling took place. Our second hypothesis, that fire frequency and extent would affect avifauna composition, with a higher abundance of savanna species in plots with higher occurrence and extent of fires, was partially supported. The results of PERMANOVA was significant for fire frequency (F = 3.56, p = 0.004; Table 3). Fire extent F = 3.23, p = 0.005; Table 3) and tree coverage F = 2.84, p = 0.007; Table 3) were also significant, but these results are not independent as they are correlated to fire frequency.

**Table 3.**
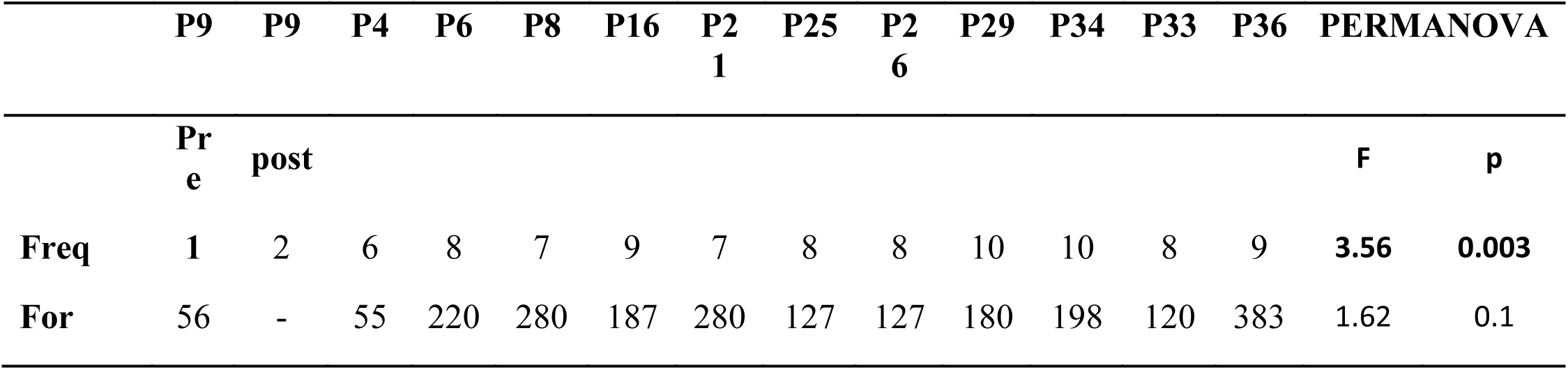
The results of F model and their respective p-value in the PERMANOVA analysis for fire frequency for the twelve years (Freq) and the distance of plot centroid for the forest (For)..

Some savanna species were not present in plots with fire-occurrence frequencies inferior to eight years, and forest species occurred mainly in the lowest extreme of the fire-frequency gradient (less than six years; Figure 4). Some forest species occurred only in plots 4 and 9, which were the plots with the lowest fire frequencies in the last 12 years. The GLM analysis indicated that savanna-species richness was not affected by any of the environmental variables we tested (Table 4). However, for savanna species abundance, fire frequency alone was the best model (Table 4). In forest species, for both richness and abundance, only distance to forest was kept in the best model (Table 4).

**Figure 3.**
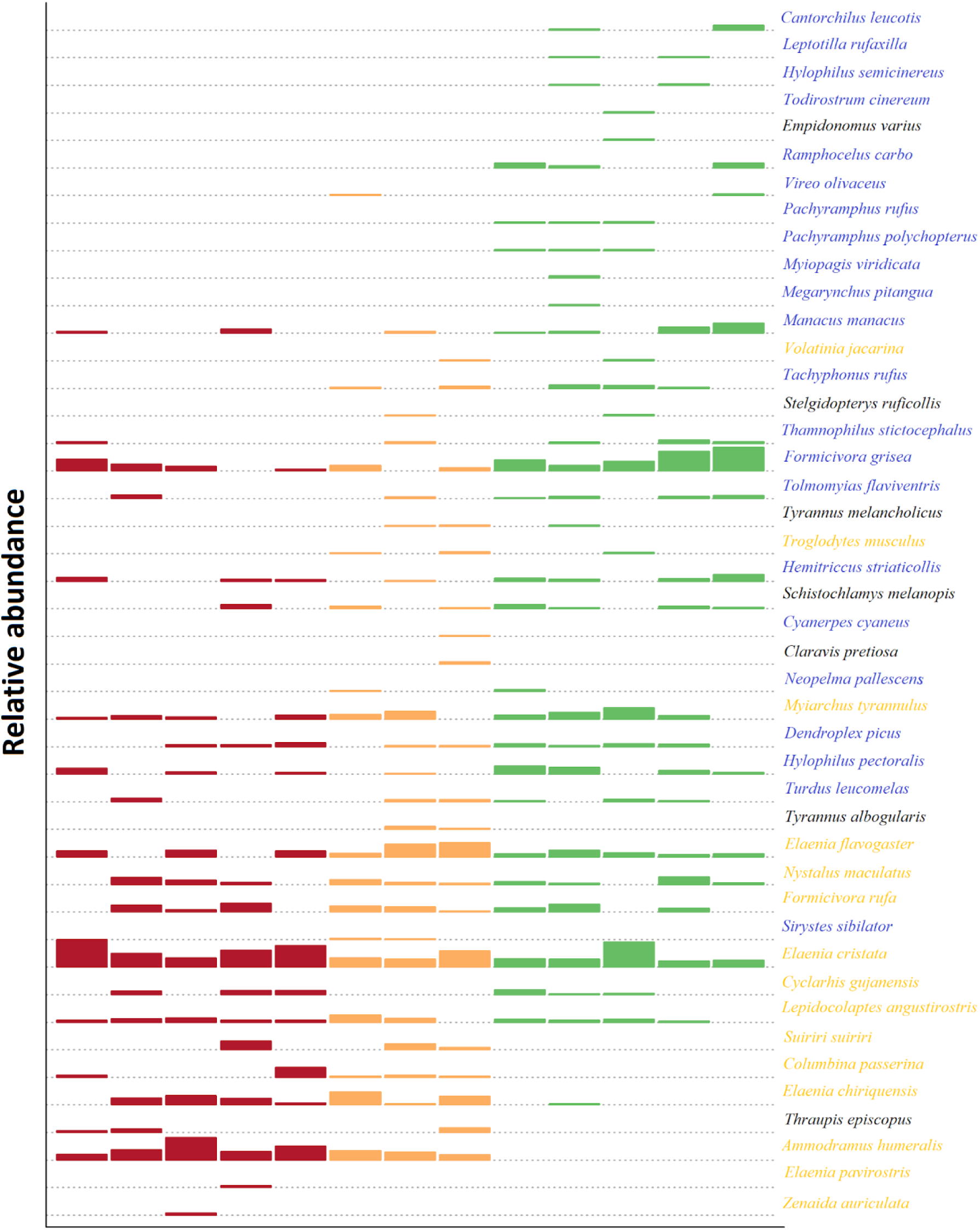
Species relative abundance in the 12 ha area of the peninsula. Each row represents a species and the bars represent the relative abundance of the species in the sample. The bar colors represent the sampling period (red = 87/88, orange = 96/97, and green = 09/10). Each column represent a sampling occasion. The samples are ordered chronologically (oldest to most recent) on the horizontal axis. The color of species name indicates that it occurred mainly in savanna (yellow), forest (blue) or the data did not show a trend regarding habitat use (black). After 13 years without fire, some savanna species disappeared of the area.

**Figure 4.**
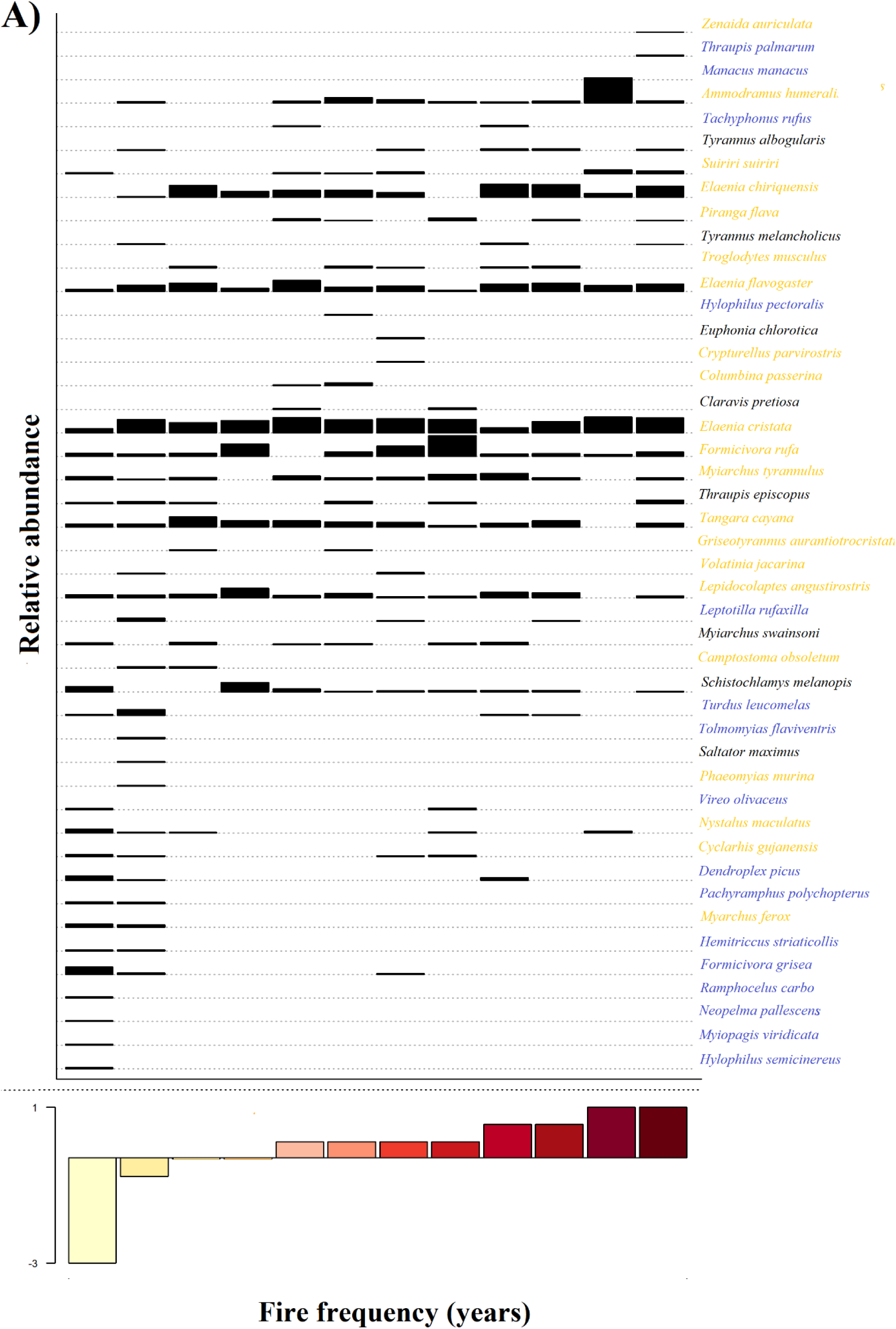
Species occurrence in the 12 savanna plots. The plots are ordered by fire frequency per year in the last 12 years. Each row represents a species and the bars represent the relative abundance of the species in the plot. The color of a species name indicates that it occurred mainly in savanna (yellow), forest (blue) or the data did not show a trend regarding habitat use (black). Some species, mostly forest species, are only present with lower fire frequency.

**Table 4.**
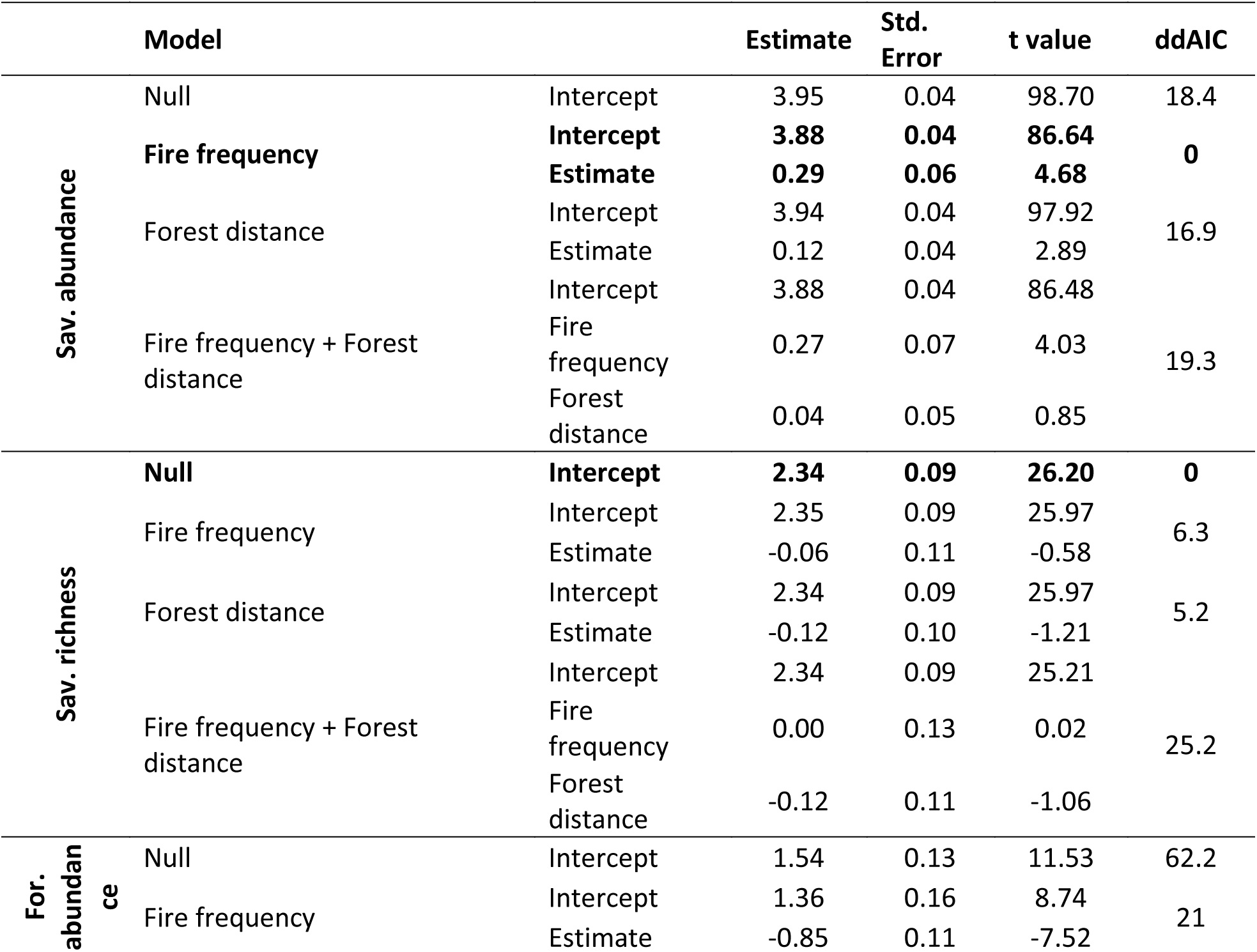
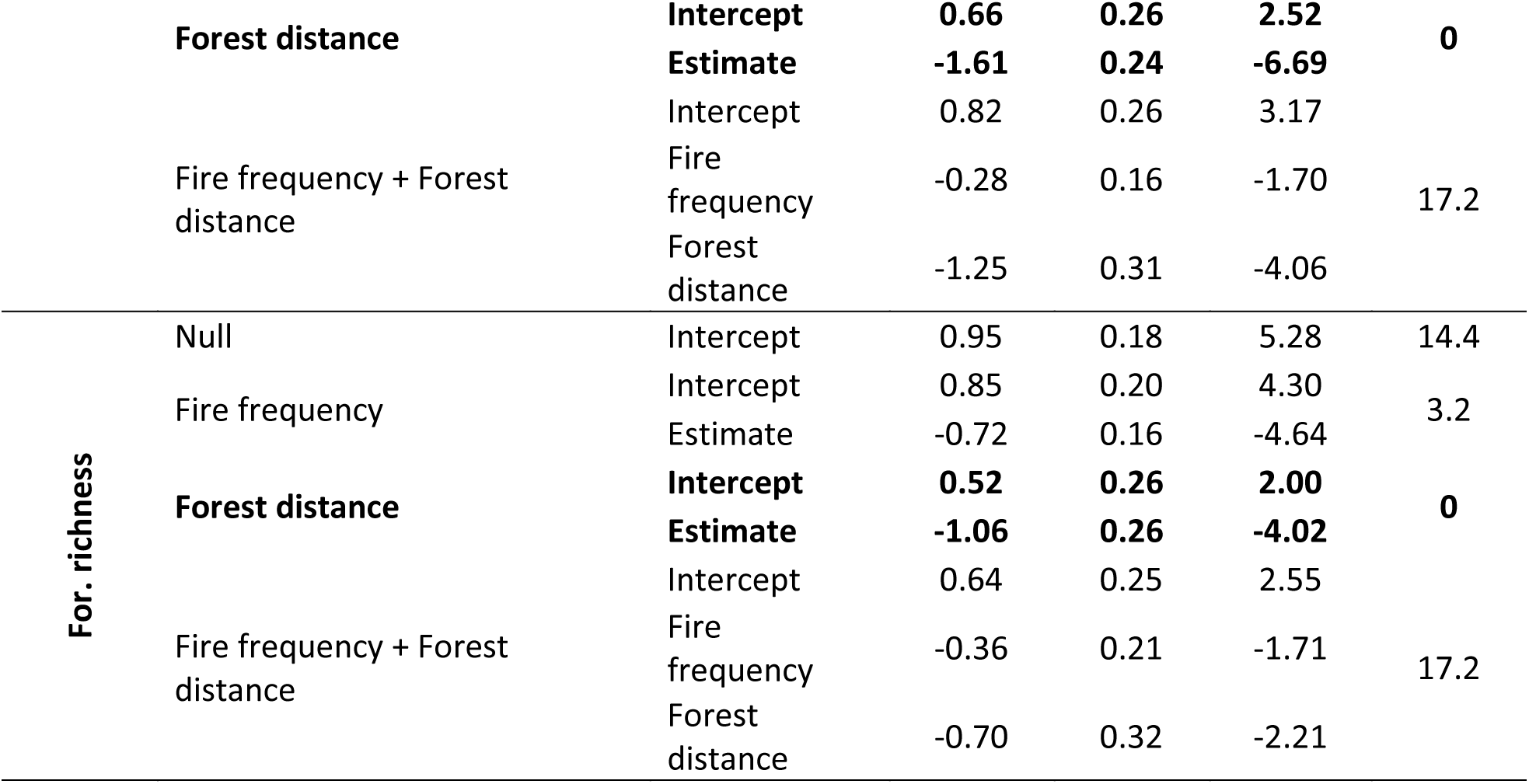
Generalized Linear Models (GLM) of savanna (Sav.) and forest (For.) Species abundance and richness with the environmental variables. Best models are in bold, for savanna species abundance the best model kept just the fire frequency while for forest species (both richness and abundance) the best model kept just the distance to the next forest.

### Additional short-term fire-effect observations

The bird species composition in plot 9 a month after the 2010 fire was different from the composition observed previously (Figure 5). Some species, such as *Schistoclamys melanopis*, *Myiarchus ferox, M. tyrannulus,* and *Dendroplex picus* were not captured after the fire, while *Claravis pretiosa* and *Ammodramus humeralis* were captured exclusively in the post-fire survey. In the long-term study, which included plot 9, these species were not captured in any of the samples in the period without fire. However, these species occurred in samples from the periods with frequent-fire regime, and *A. humeralis* was one of the most frequent species in those samples (Figure 3).

**Figure 5.**
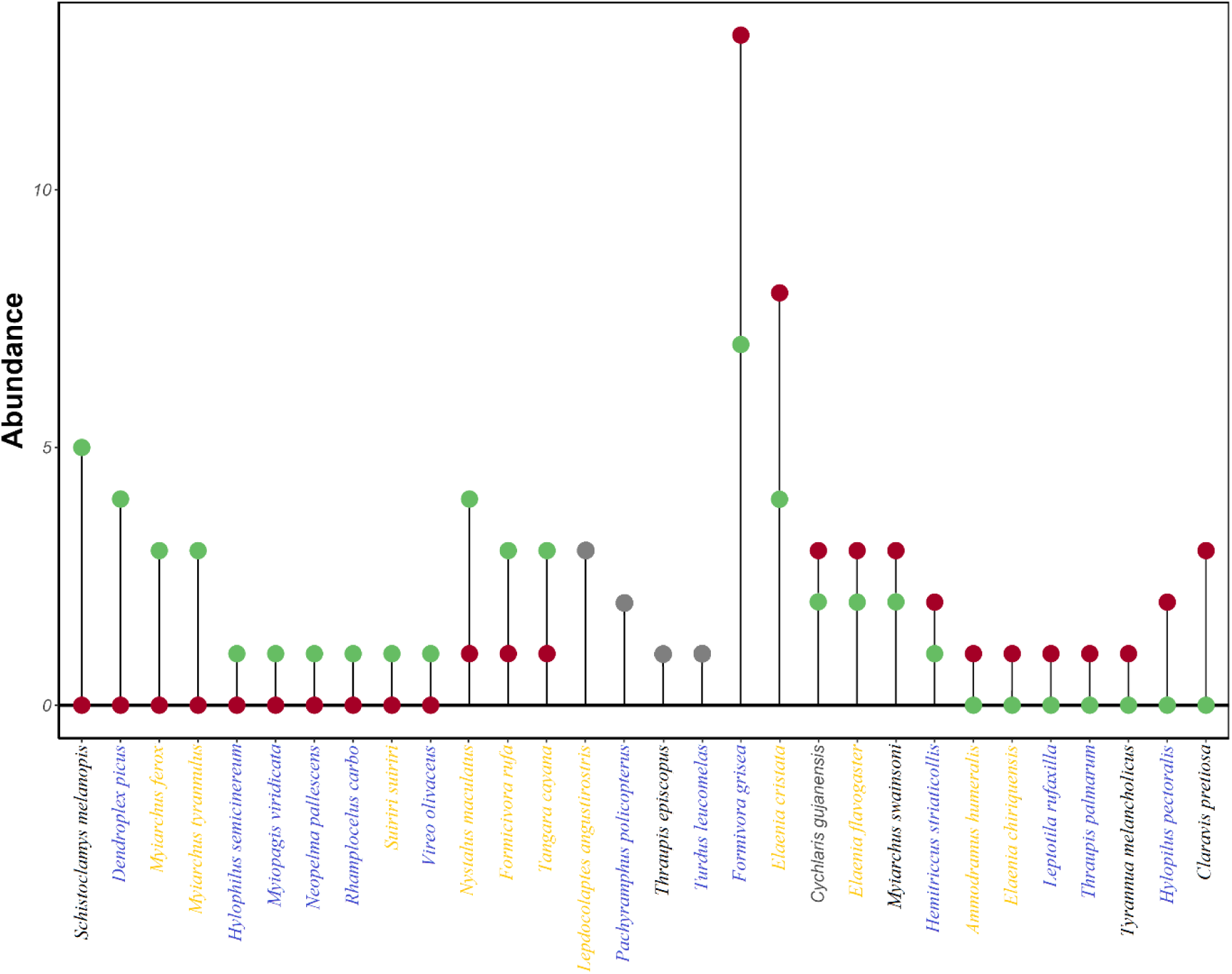
Changes in the bird community in plot 9 before (12 years of fire suppression) and after the fire in 2010. The y-axis represents the difference in abundance of each species captured pre-fire (green dots) and post-fire (red dots). For species with the same abundance pre- and post-fire the dots are grey. The color of a species name indicates that it occurred mainly in savanna (yellow), forest (blue) or the data did not show a trend regarding habitat use (black). Ten species were captured only in pre-fire period, while seven were captured only in the post-fire period.

## Discussion

We evaluated the temporal and spatial change in bird assemblages under distinct fire regimes in an Amazonian savanna. Although we had only three periods with a total of 13 samples (five in 1988/89, three in 1997/98, and five in 2009/10 after 13 years without fire), the avifauna composition included more forest species. Similarly, even though only one time period was sampled in the spatial analysis, with just one plot that did not burn, fire frequency affected the avifauna, with some savanna species appearing soon after a fire in a plot that had not previously burned. Although it is not possible to make statistical comparisons for the short-term observations, they were consistent with those in the longer-term studies, with some savanna species appearing soon after a fire in that plot. Each part of the study should be analysed with caution due to limited sampling, but the temporal and spatial analyses reinforce each other and indicate the importance of fires for savanna species.

### Temporal analysis

The dynamics of local bird assemblages in our study changed over time in accordance with changes in fire occurrence. Regular fires maintained the area suitable for an avifauna comprising typical savanna species, but periods without fire led to an influx and consequent predominance of forest species in the bird assemblages, as found for birds and other vertebrates in an Australian savanna under fire suppression for 23 years (Woinarski et al. 2004).

During the period without fires, there was a marked increase in woody-vegetation cover, mainly due to an increase in density of tree species associated with forests in the study region (Lima et al. 2020). Forest birds foraging in adjacent savannas potentially disperse forest seeds into the savanna (Tubelis 2004). Therefore, it is likely that the influx of forest birds has intensified the process of forest-tree recruitment by seed dispersal in the savanna area. The main factor that restricts the establishment of forest-tree seeds in savannas is fire, in spite of seed morphological constraints, thus, fire suppression favours the expansion of forest species into the savanna (Hoffmann et al. 2004, de Souza Campos and Jardim 2020, Lima et al. 2020), and promotes woody-cover increase.

Shrub encroachment (*i.e.* woody-cover increase) has been indicated as a driver of changes in bird assemblages in African savannas, especially due to the loss of species associated with open savannas (Krook et al. 2007, Sirami et al. 2009, Sirami and Monadjem 2012). In our samples, two species that were among the most frequent species from 1987 to 1997, *A. humeralis* and *C. passerine*, were not recorded in the study site in periods without fire. These species are ground dwellers and probably associated with grass cover, a characteristic ubiquitous in African savanna birds that are undergoing habitat loss due to shrub encroachment (Sirami and Monadjem 2012, Abreu et al. 2017). Savannas protected from fire have less grass cover and denser woody vegetation in relation to savannas undergoing frequent burning (Moreira 2000, Woinarski et al. 2004, Passos et al. 2018). The passage of fire reduces the density of the middle stratum of the vegetation and reduces shrub-cover extent (Sanaiotti and Magnusson 1995), favours the expansion of grasses (Moreira 2000, Fidelis and Zirondi 2021), impairs access of birds to the soil (Crawford 1979), and it is likely that a similar process occurred in our study site. In the savanna we studied, fires occur during the late dry season, which coincides with the nesting season of many species of savanna birds (Sanaiotti and Magnusson 1995).

Savanna birds have high specificity in terms of vegetation structure (Tubelis and Cavalcanti 2001, Krook et al. 2007, Sirami et al. 2009) and the direct effects of fire on birds is expected to be less pronounced than the strong response of birds to changes in the environment. The interaction between the direct and indirect effects of fires on the biota makes it difficult to consider them separately. We did not detect direct short-term effects of fire on individual birds. In contrast, many individuals were recaptured after fires in the short and long-term, a result also observed previously in the same region (Cintra and Sanaiotti 2005) and in the Cerrado of Central Brazil (Piratelli and Blake 2006).

### Spatial analysis

The effects of fire frequency, fire extent, and tree cover were not distinguishable due to their high covariance. Fire inhibits the establishment of young trees and sprouts, while adult trees in savannas survive burns (Prior et al. 2010). In Amazonian forest areas, fires are more frequent and cover larger areas in deforested areas (Sorrensen 2009; Lima et al. 2012), due to the decrease of humidity (Silvestrini et al. 2011). The high correlation between fire frequency and fire extent in our study area probably results from a similar process. The fires suppress the vegetation making the area more susceptible to future fires that can burn a bigger area compared with unburned regions. We expected that plots under a regular fire regime would have little variation in tree cover over time because of the inhibition of tree recruitment by fire, and consequently it would be the vegetation strata least correlated with fire history. Therefore, we chose tree cover as the main vegetation variable. However, tree cover was highly correlated with fire history, impairing the distinction between the effects of these two variables (Camill et al. 2009).

A fire-severity index that combined fire extent, shrub-cover loss and branch diameter (a proxy for fire intensity) affected birds in European mountain rangelands (Pons and Clavero 2010). Since fire regime is a multifactorial disturbance, which probably has a multitude of synergistic direct and indirect effects on the biota, it is challenging to assign organism responses to particular aspects of a fire regime. Here we used fire frequency, as the proxy of fire regime since it covaries with fire extent and tree coverage.

Some species occurred almost exclusively in the extreme of the fire frequency and tree-cover gradient. Plots 4 and 9 had a similar and extensive tree cover, but the fire frequency in plot 4 was pronouncedly higher. The effect of the interaction between these variables, although very plot-specific, is illustrated by the occurrence pattern of some species in plot 4. Even though plot 4 shared some forest species exclusively with plot 9 (e.g. *Hemitriccus striaticollis* and *Pathyramphus poluchopterus*), some savanna species (*e.g. A. humeralis* and *E. chiriquensis*) that occurred in plot 4 were only captured in plot 9 after a fire. In spite of its extensive woody vegetation, its fire frequency probably also favoured the occurrence of some typical savanna species.

The spatial variation in the fire regime, may contribute to the bird diversity at the landscape level. As tropical forests and savannas seem to represent alternative states in some areas, such as the Amazonia where savannas suppressed of fires tend to have more forest species present (Magnusson et al. 2002). The forest-savanna transition may be associated with thresholds or tipping points in environmental variables, such as fire frequency (Hirota et al. 2011, Lehmann et al. 2014). Tropical forest burned several times became a savanna and savannas surround by forests, after long periods without fire became more forested with typical forest species returning. As this transition is dynamic some species could occupy one or another. Indeed, in another Amazonian savanna landscape, most bird species crossed from one habitat to another (de Sousa et al., 2022), and in our study distance to the forest was the main factor explaining forest species richness and abundance.

### Implications for conservation

Amazonian savannas share species in common with the Cerrado, allowing the expansion of the Central South America savanna biota to the region (Aleixo and Poletto 2007, Ritter et al. 2020), including some threatened species, such as *Cercomacra carbonaria* and *Synallaxis kollari* (Santos and Silva 2007). Amazonian savannas are important for conservation at both local and regional levels, since they occur surrounded by the tropical forest, forming mosaics of complex vegetation (Adeney et al. 2016), which increase the beta diversity. Furthermore, Amazonian savannas shelter several Cerrado species with distinct evolutionary and biogeographic characteristics (Garzón-Orduña et al. 2014, Resende-Moreira et al. 2019, Ritter et al. 2020).

Savannas are generally distributed in areas with low annual precipitation and high precipitation seasonality (Lehmann et al. 2011). However, many tropical savannas currently occur in areas that are predicted to support forests (Bond and Keeley 2005, Murphy and Bowman 2012). In these areas, disturbances, such as fires, help to keep the vegetation open (Fidelis and Zirondi 2021). Fire histories that favoured forest species and culminated in a higher richness were in general detrimental for savanna species (Durigan 2020). However, a frequent-fire regime may lead to species loss and ecological disequilibrium (Medeiros and Miranda 2005, Fidelis and Pivello 2011). In ecosystems, such as the Cerrado, where patches of forest harbour a great portion of the diversity (Silva 1995), it is advantageous to apply comprehensive conservation strategies that include less-frequent fire regimes. However, Amazonian savannas have a limited extent, are under-protected and threatened, and efforts to conserve their diversity should be concentrated on the savanna species strongly associated with this habitat. Indeed, a study focusing on forest habitats in the same area of this study showed that small forest fragments can maintain a large portion of bird communities (80%) of the continuous-forest area (Cintra, Magnusson and Albernaz, 2013), highlighting the higher vulnerability of savanna species birds in the region.

The savannas of Alter do Chão are all on private property, resulting in an idiosyncratic and unplanned fire regime. This allowed the existence of a dynamic community over time and space. Regrettably, areas of tropical savannas are under pressure for agriculture and livestock grazing, resulting in an alarming rate of degradation. Recent changes in Brazilian environmental policy have resulted in an increase in the rate of deforestation in Amazonia (Pereira et al. 2019, Rapozo 2021), many times associated with fires (Silva et al. 2021). Although fire is a key factor for the maintenance of typical savanna species, the increase of anthropogenic fires associated with climate change, which increases the frequency and severity of Amazonian droughts, amplifying fire frequency, extent, and intensity (Xu et al. 2020)(Xu et al. 2020(Xu et al. 2020, may lead to the loss of bird diversity. Hence, avoiding the largest uncontrolled and illegal fires is important, but keeping small-scale fires may be a better strategy than trying to avoid fires altogether or manage complex and costly fire regimes (Andersen and Hoffmann 2011).

One of the main attractions for tourists in the region are the savannas, which do not occur in most parts of Amazonia. The local people interested in tourism could maintain the savannas, and their bird fauna, by controlled burns at widely spaced intervals that are not coincident among savanna patches, but this would require consultation with a wide variety of landowners.

## Conclusion

Fire affected bird assemblage composition at several temporal and spatial scales, and the effects were congruent. Areas under frequent fire disturbance had a predominance of savanna species in relation to forest species, while some forest species were more frequent in areas with lower fire frequency. The congruence of the results highlights validity of space for time substitution when temporal data are unavailable. However, the use of these two scales provides complementary information and allows a more complete comprehension of the effects of fire on the fauna. Fires are a key element in maintaining the eastern-Amazonian savanna-bird communities, but climate change and increased frequency and intensity of anthropogenic fires may threaten bird diversity if the subtle effects are not understood.

## Acknowledgments

We thank the Nobre brothers for support and dedication to the project, Mario Cohn-Haft for help in species identification and Paulo E.S. Massoca for the help with the production and analysis of maps. WEM received a productivity scholarship from CNPq.

## Declarations

### Ethical Approval

No ethical requirement was necessary. Birds captured in this study were marked with metal bands provided by CEMAVE-ICMBio (authorization number 1177 / 4, ICMBIO / SNA) or coloured plastic bands.

### Author contributions

L.A.C. e T.M.S. conceived the study; L.A.C., A.P.L., R.C., W.E.M. e T.M.S. collected data. L.A.C. and C.D.R. analyzed data and wrote the manuscript with contribution of all authors.

### Availability of data and materials

The html with the scripts will be provided as supplementary material and the script and tables used here are available at mega.

---

**Table S1:**
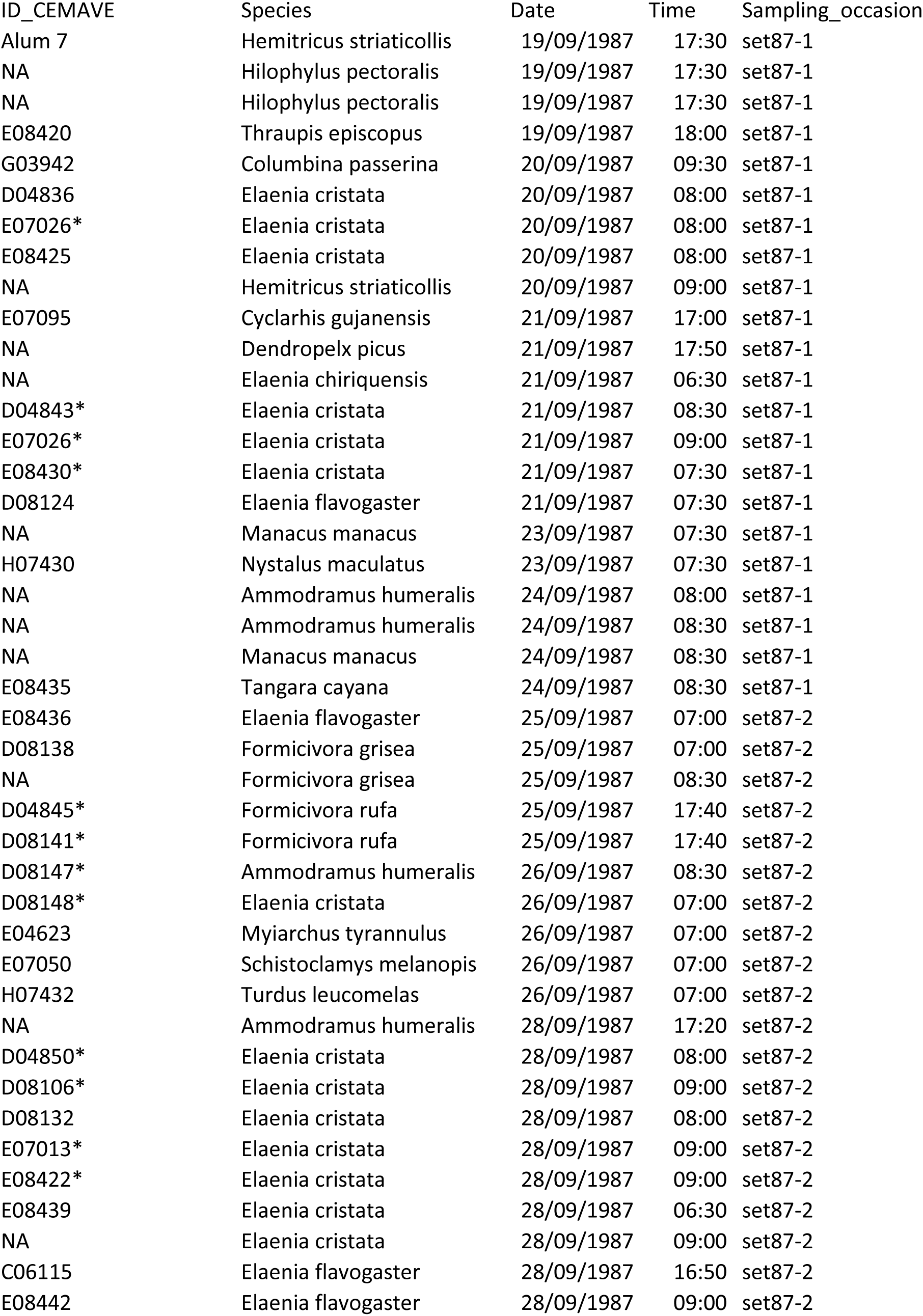
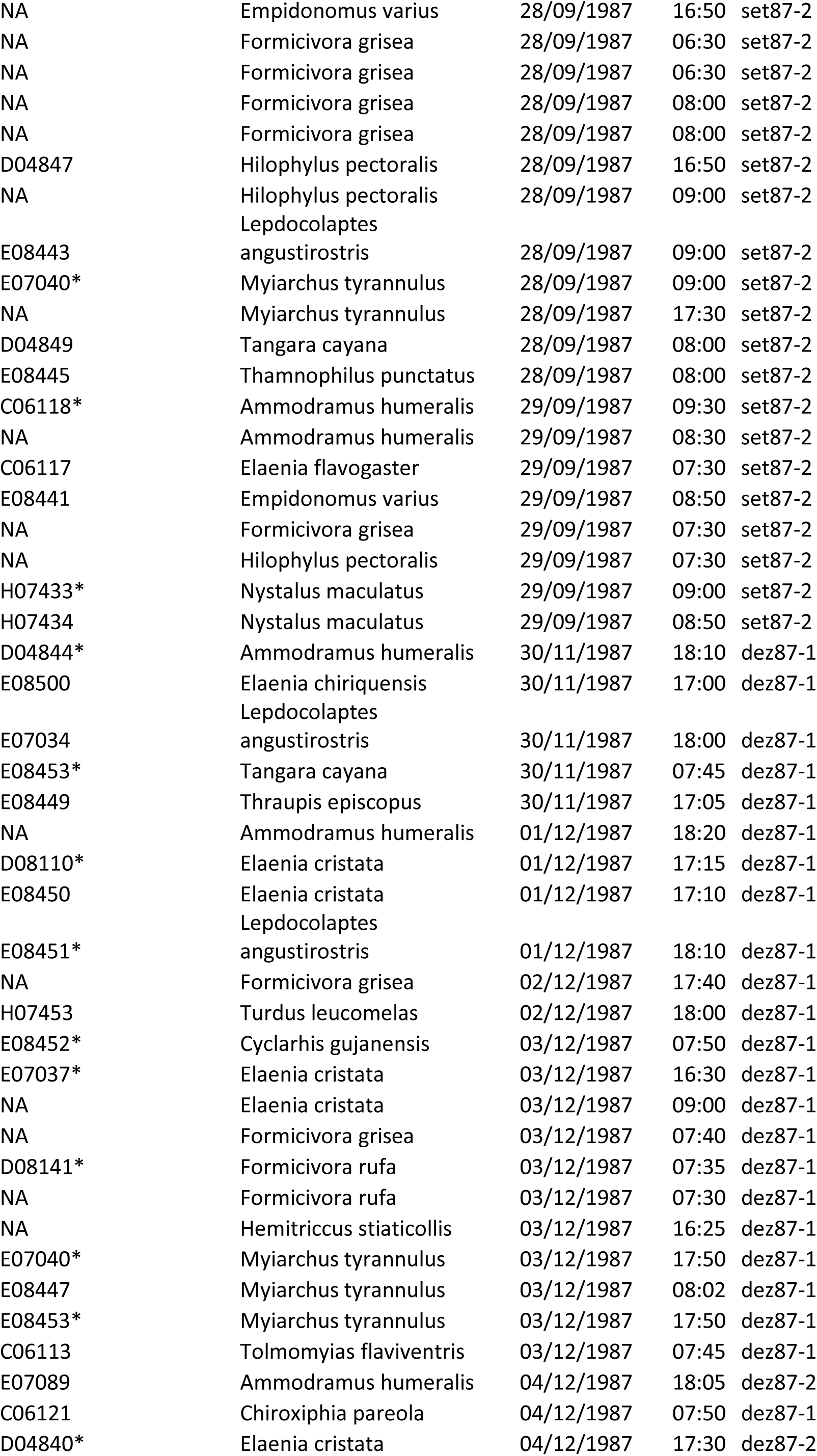
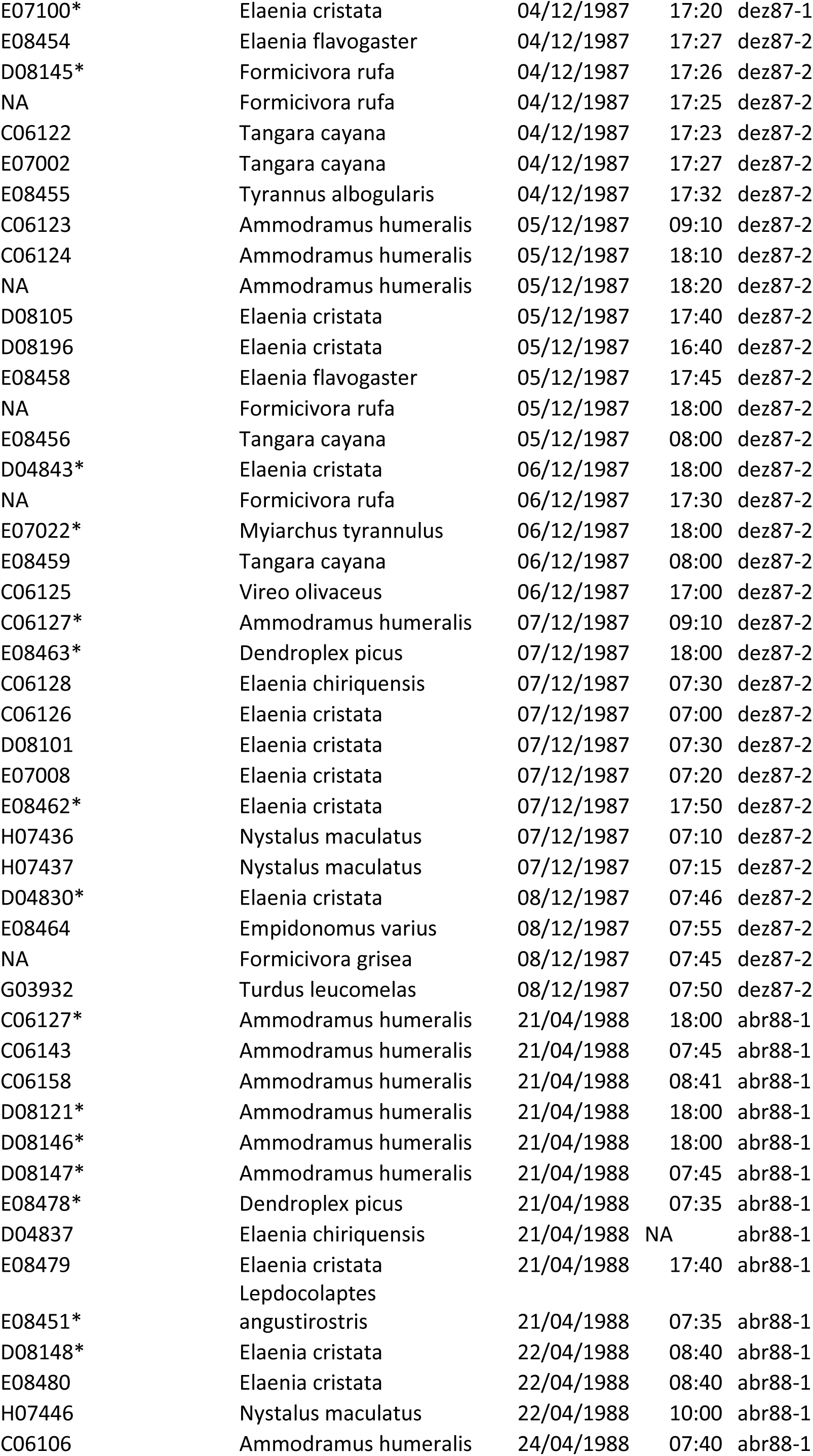
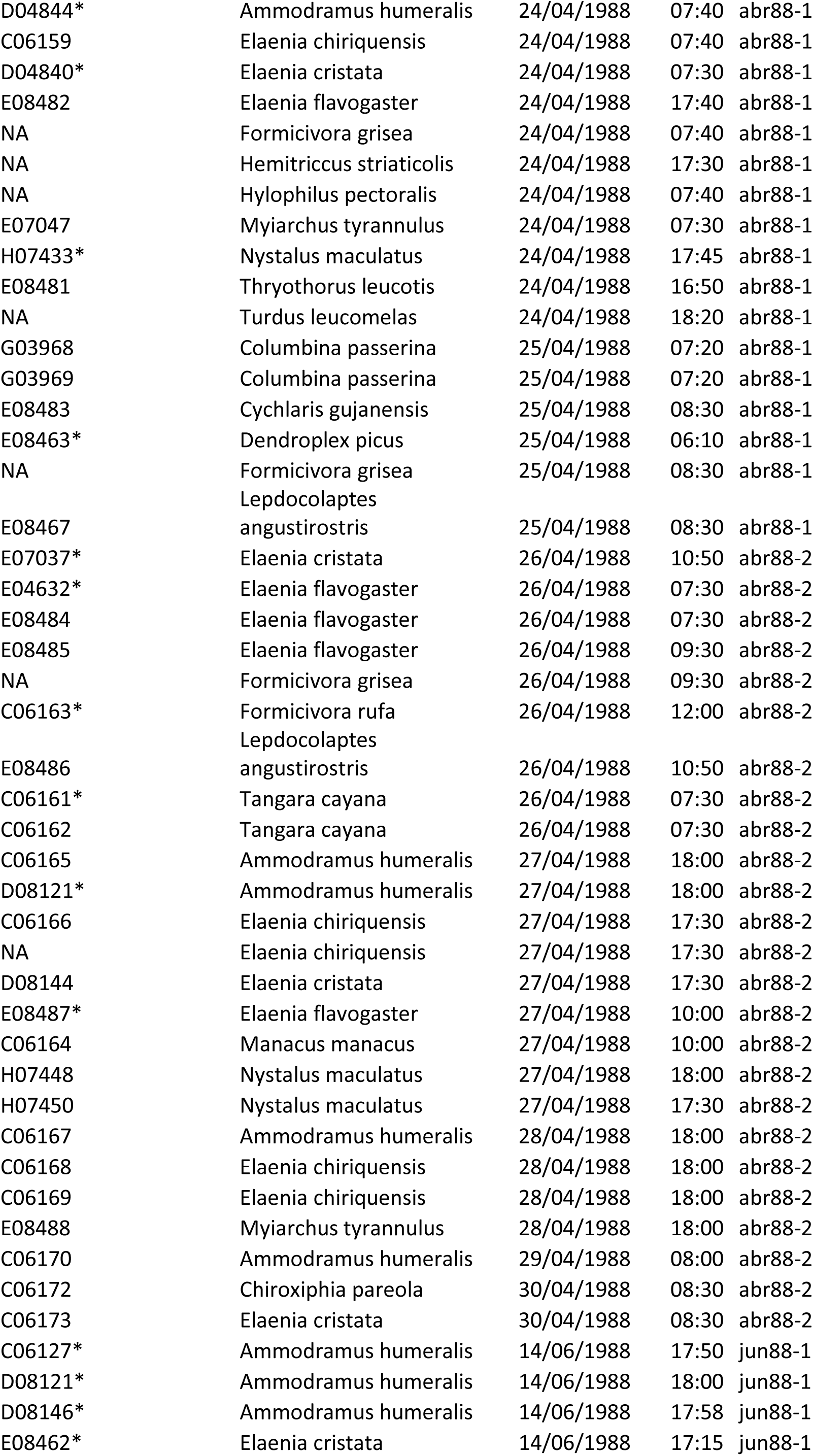
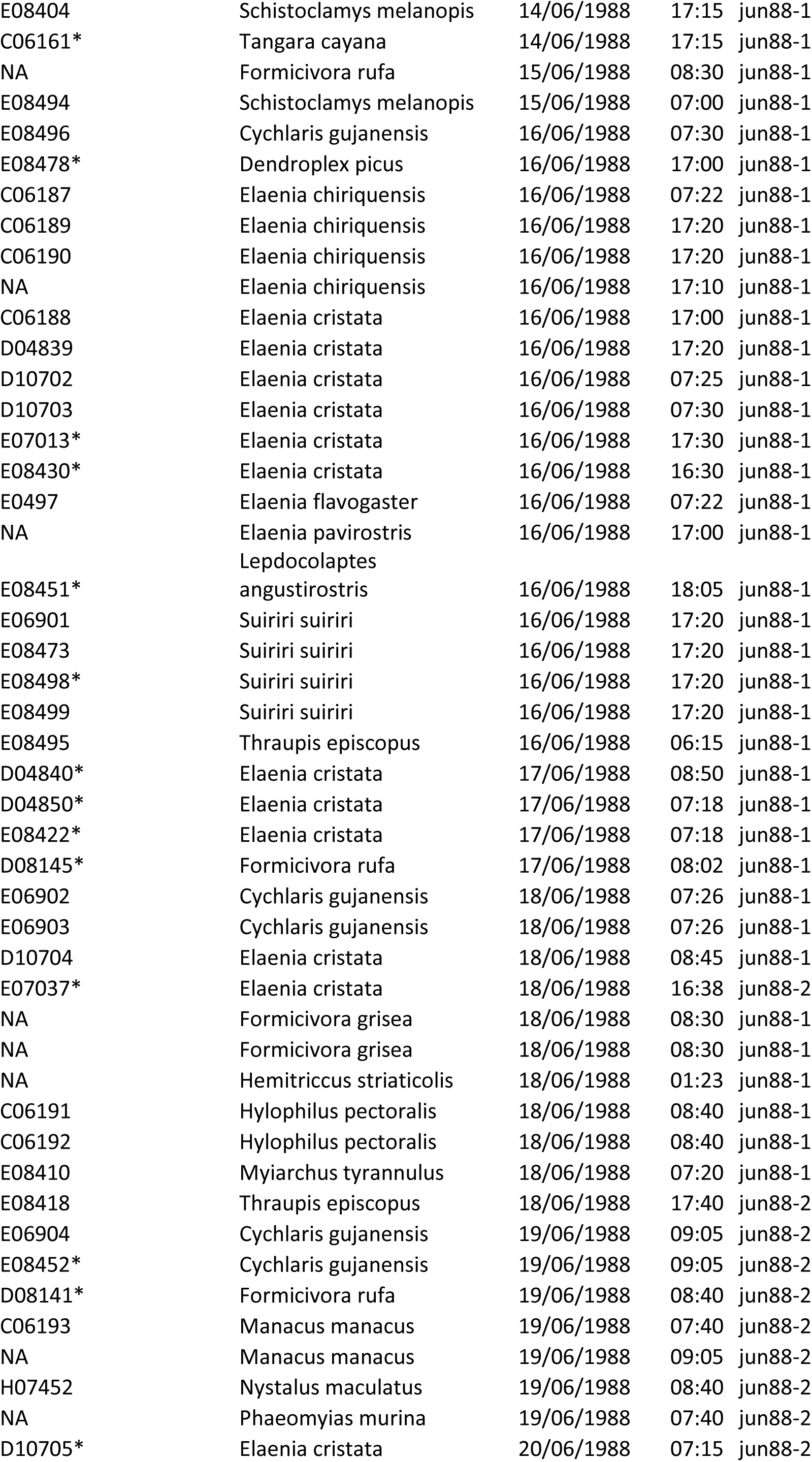
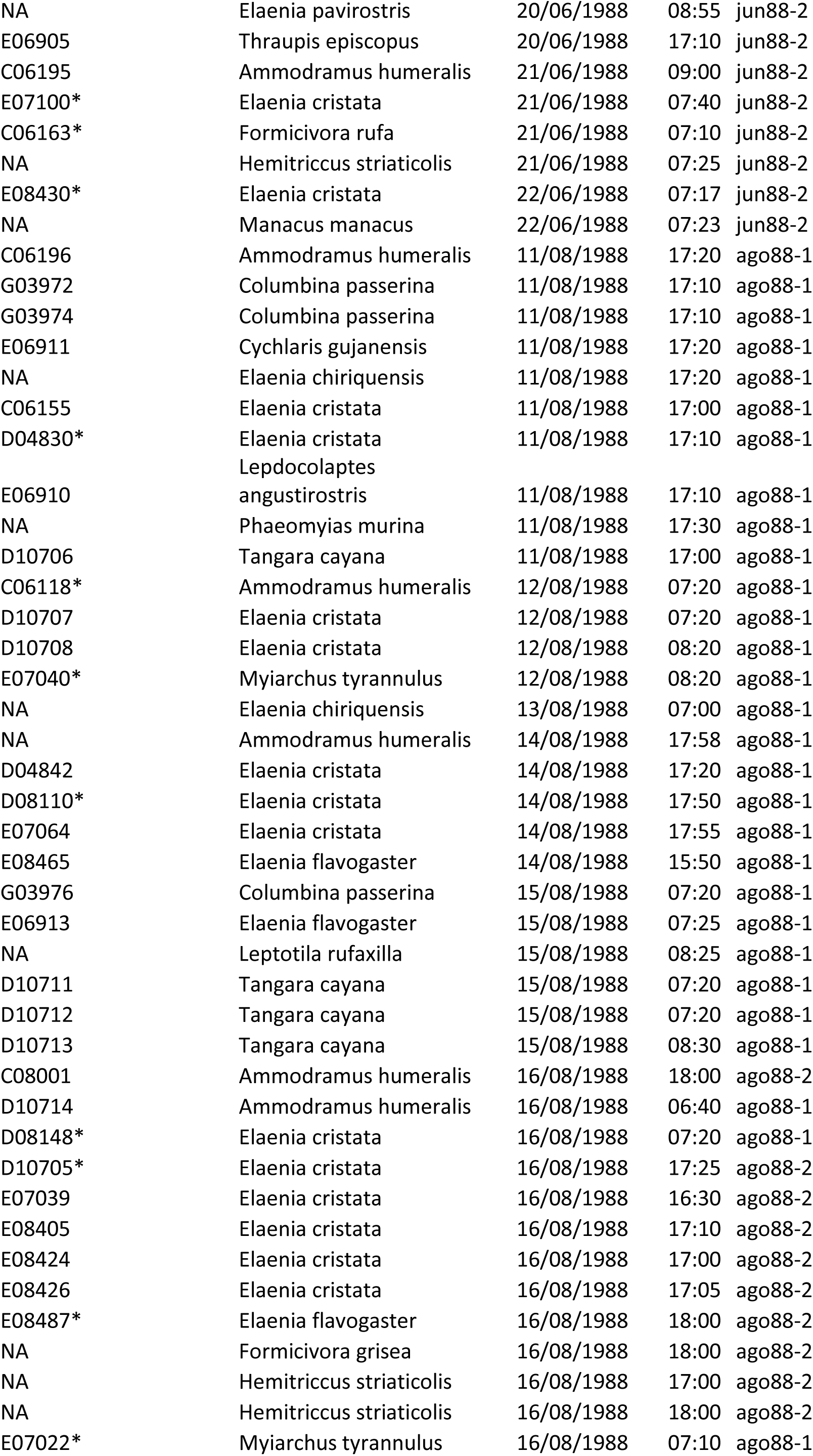
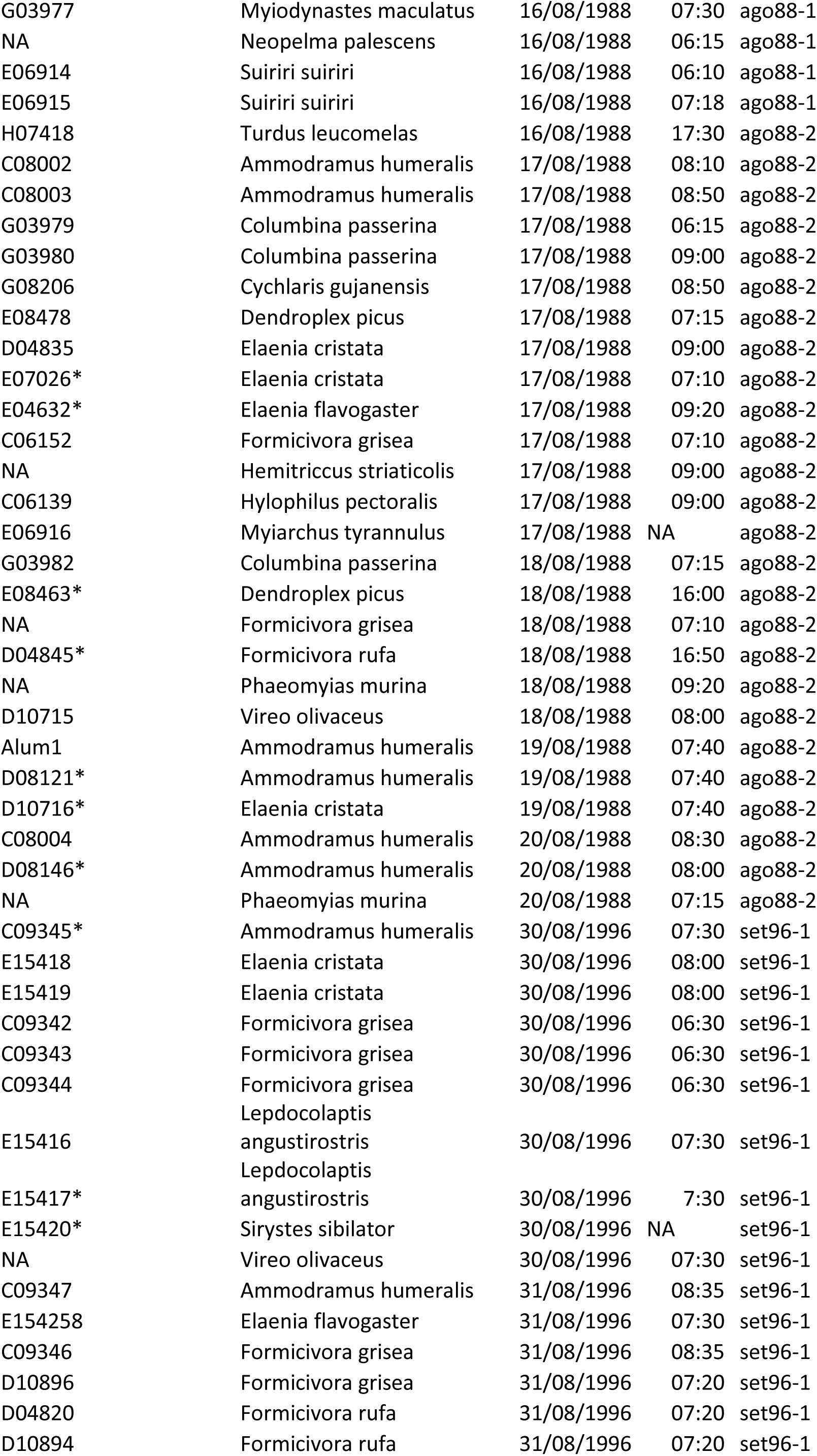
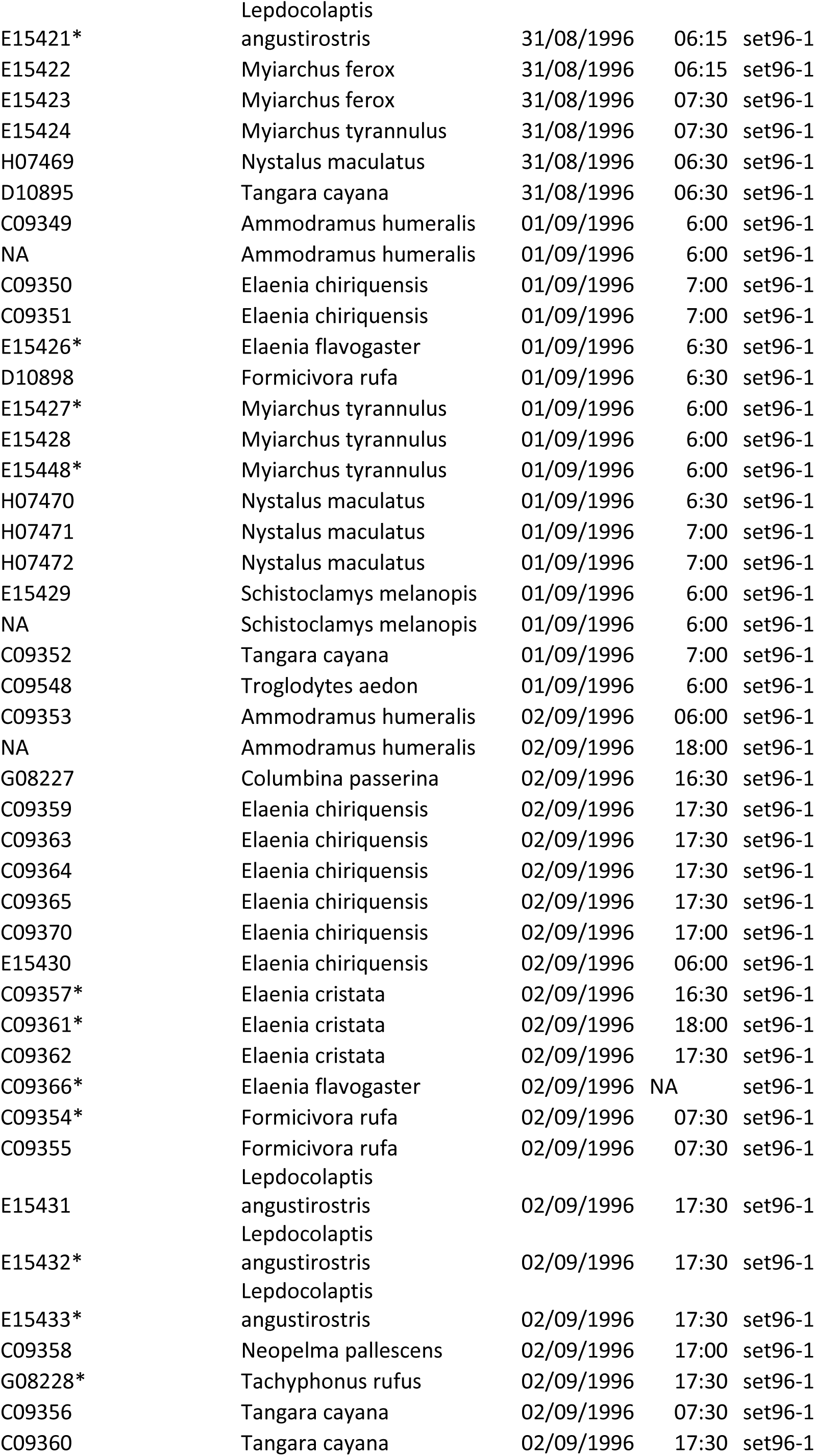
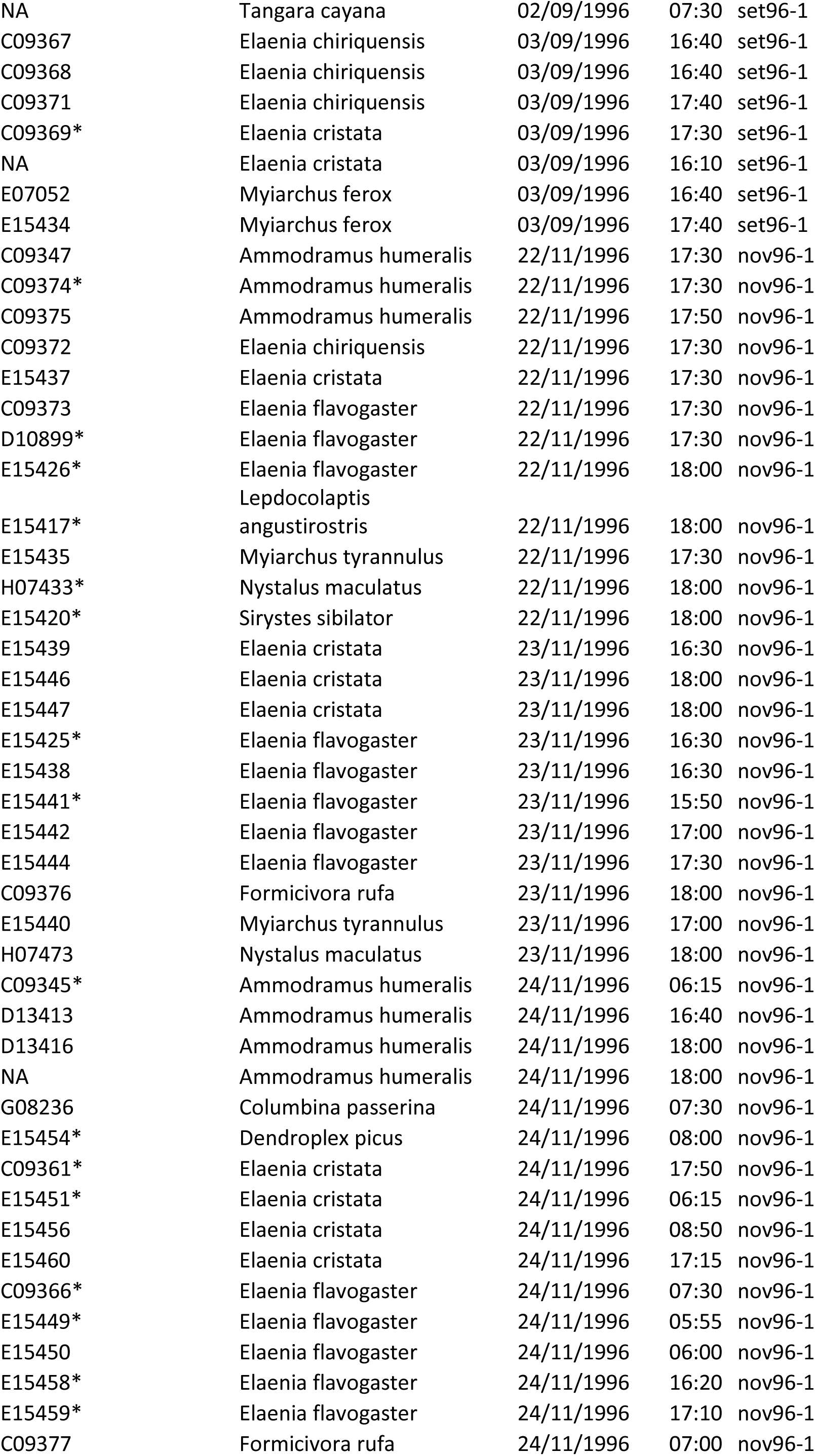
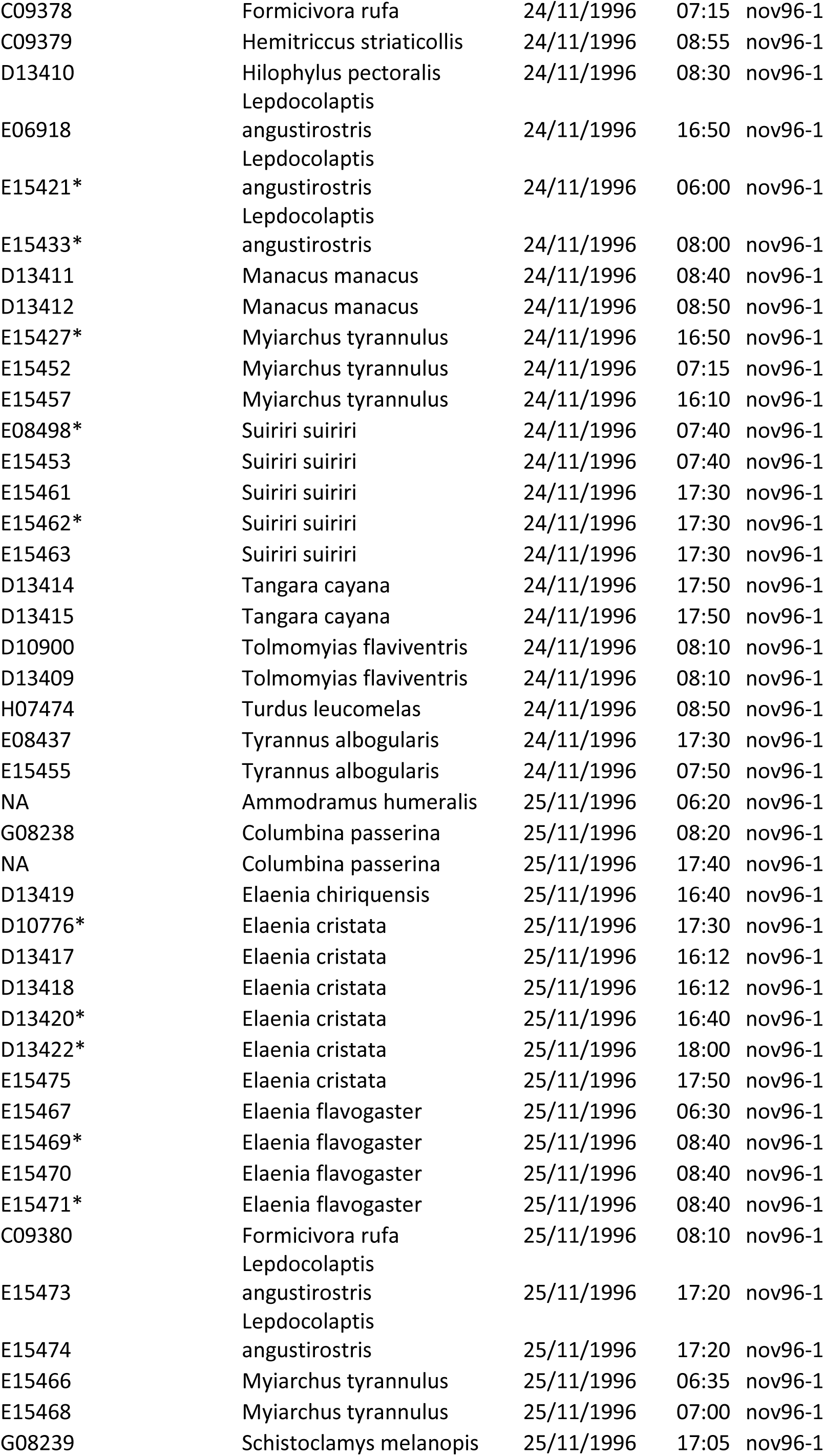
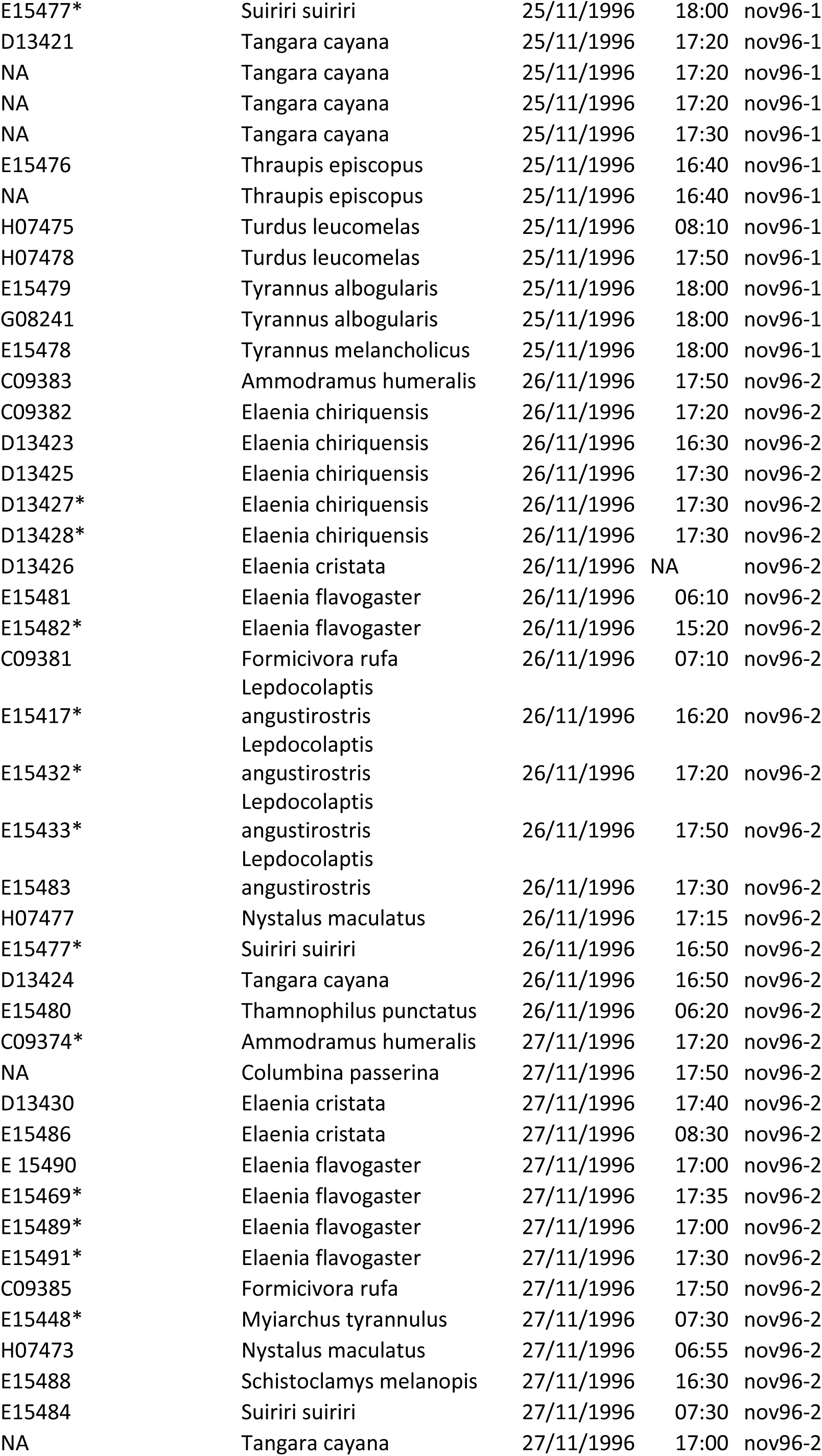
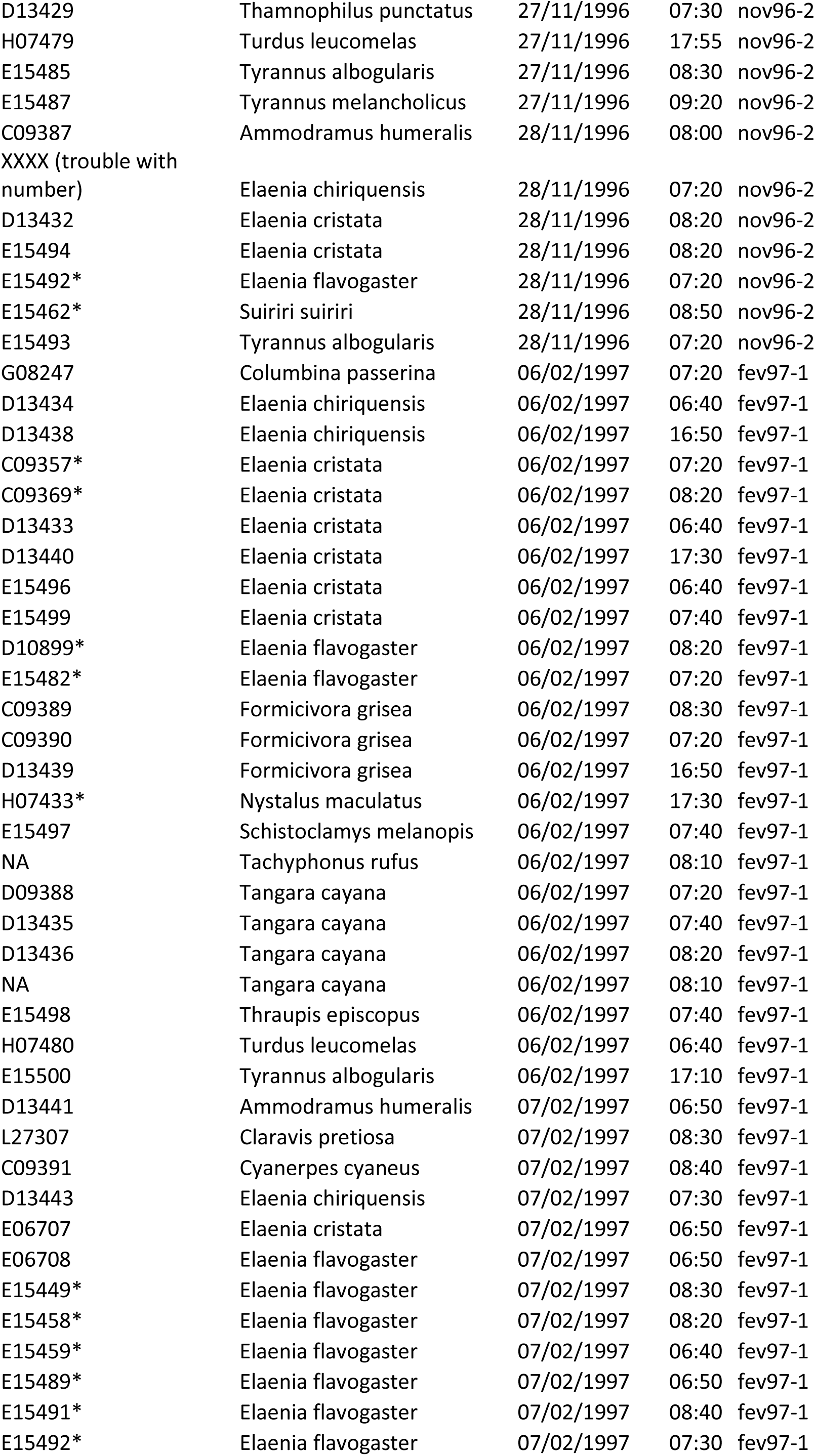
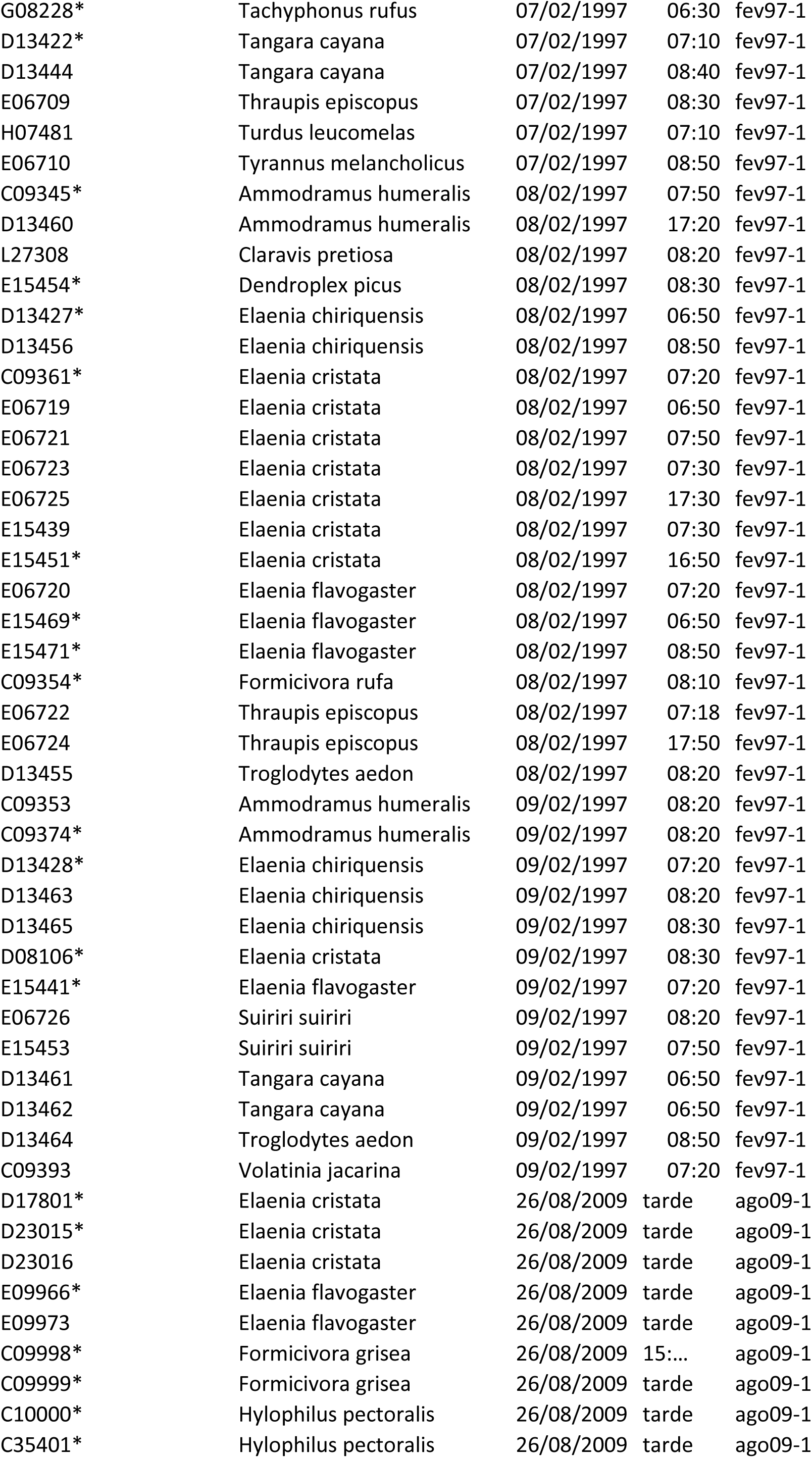
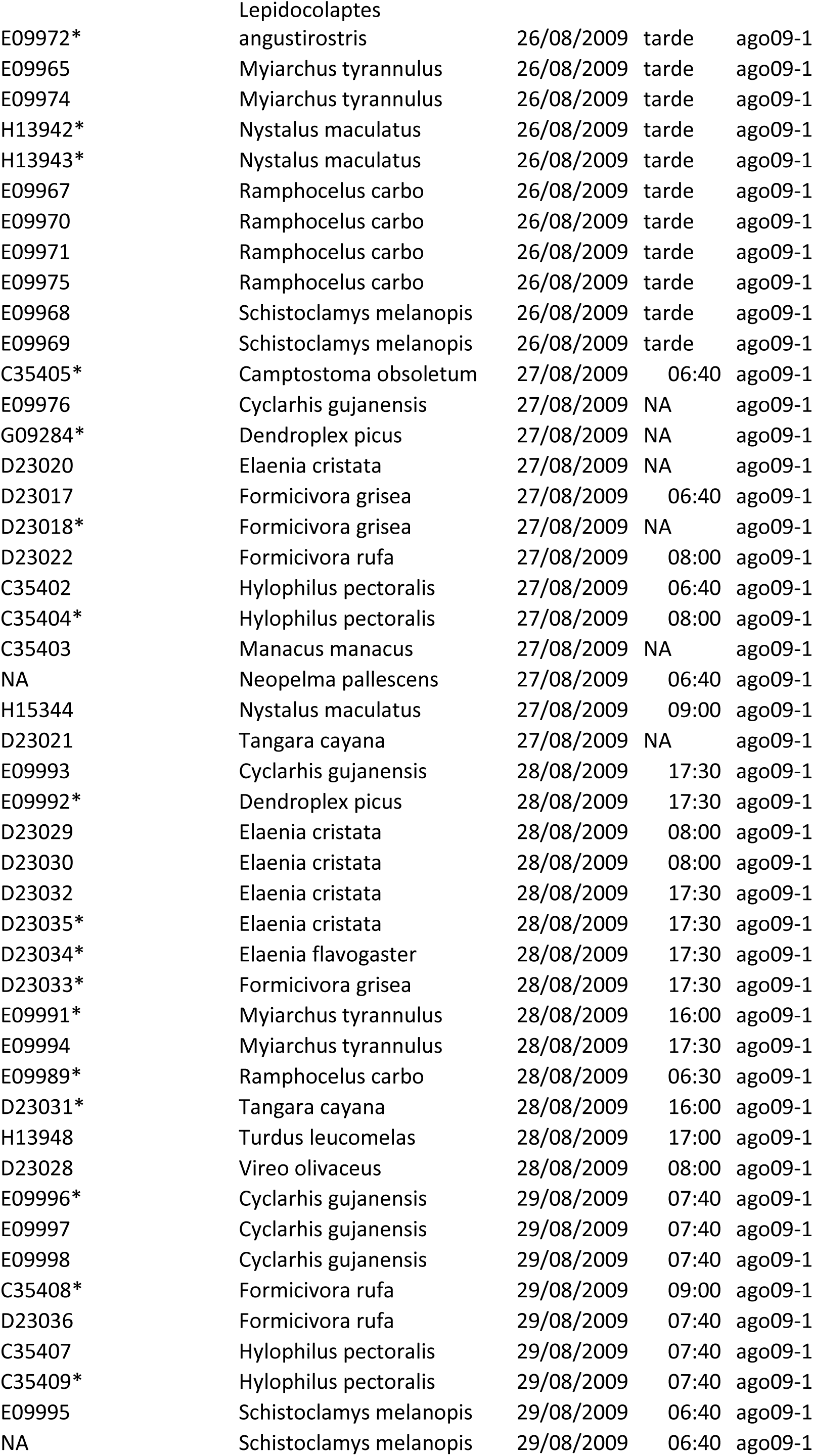
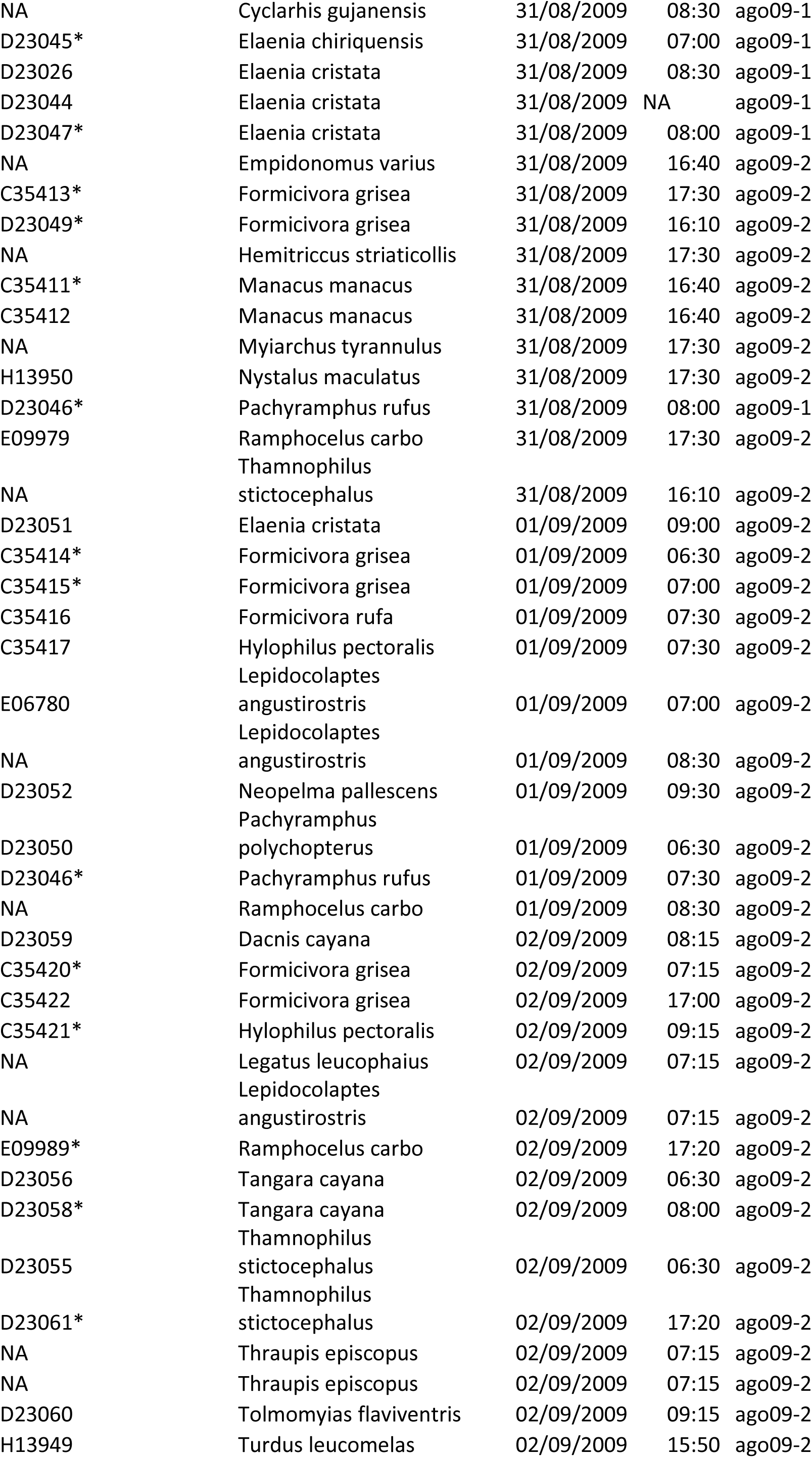
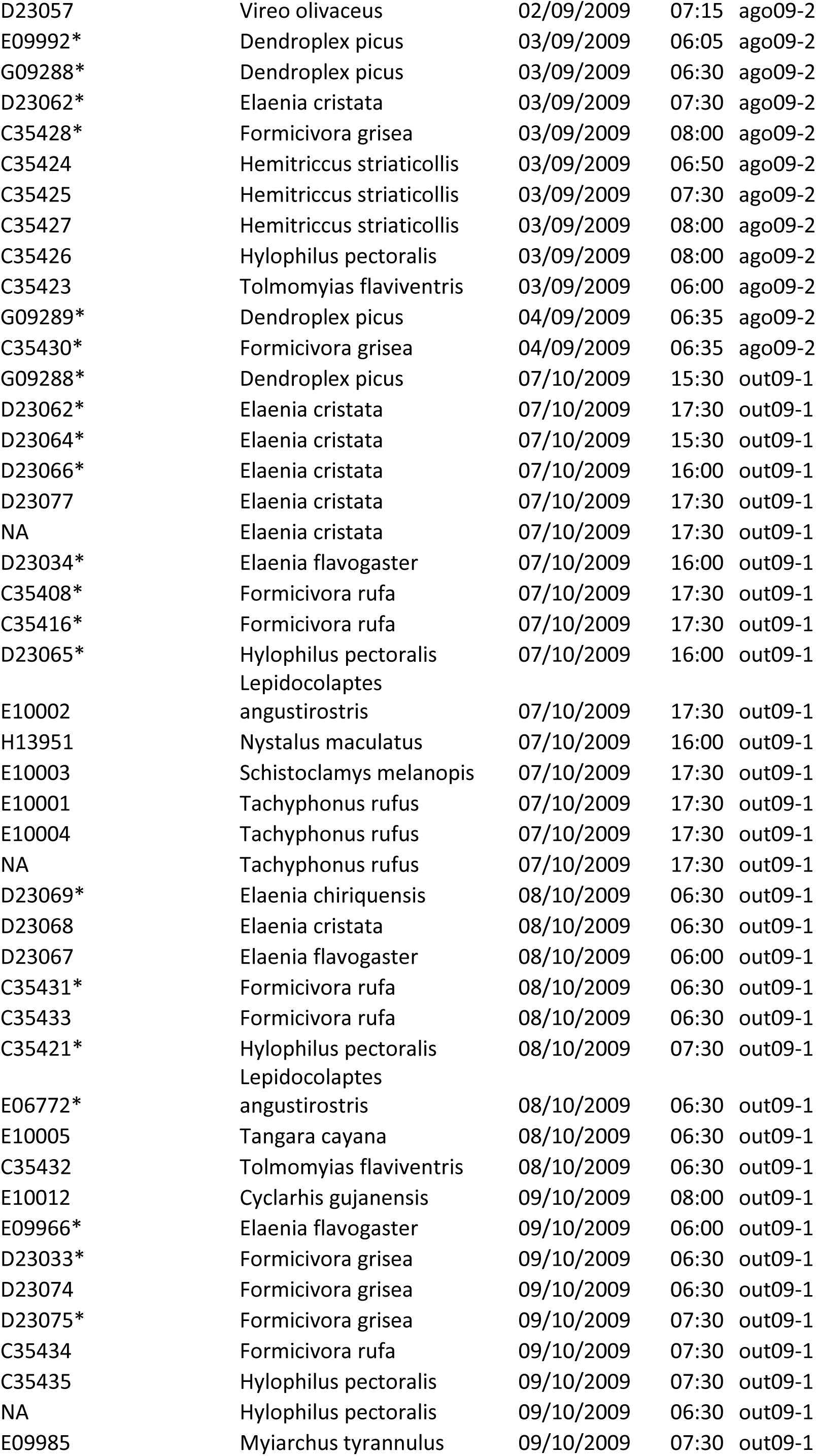
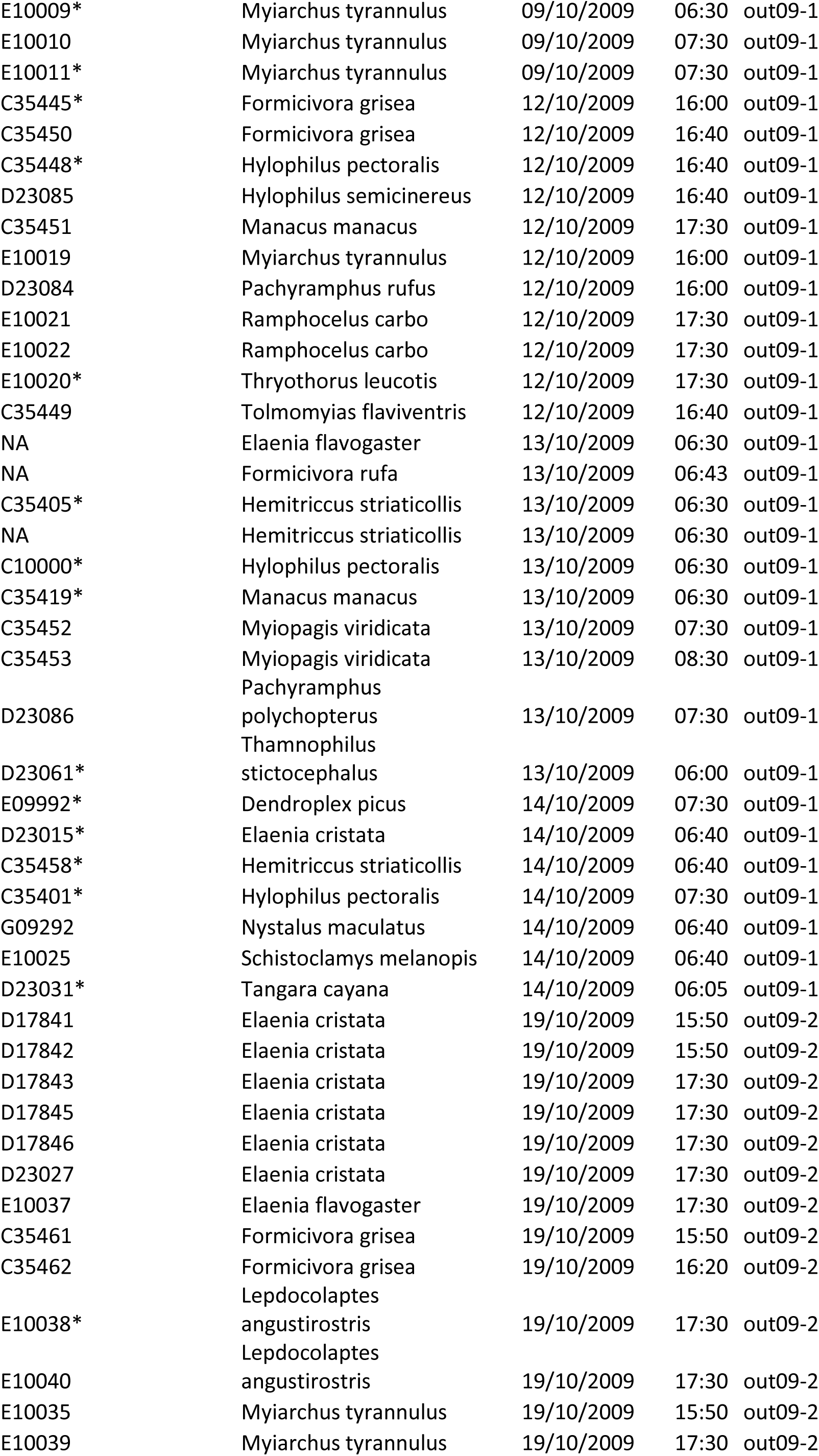
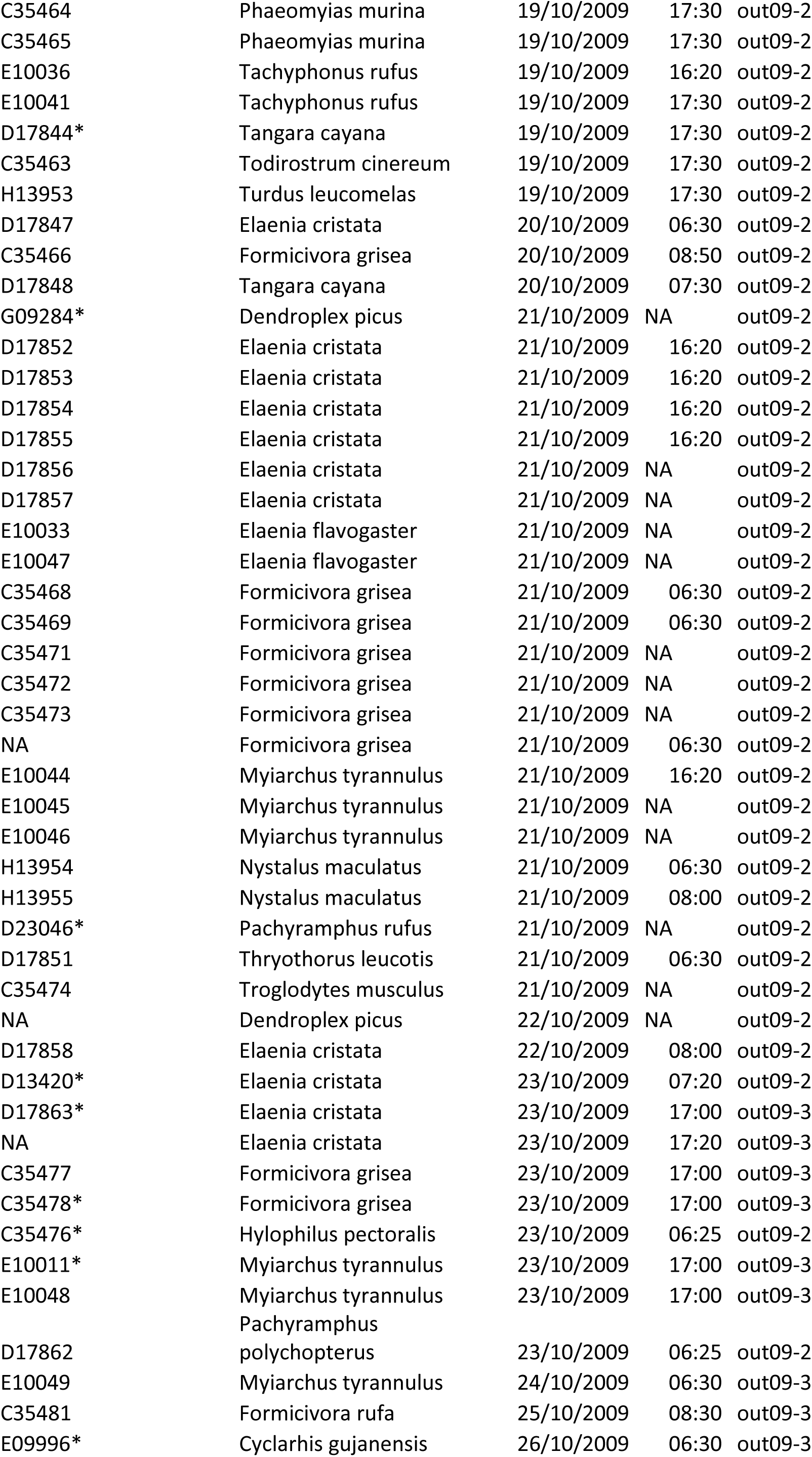
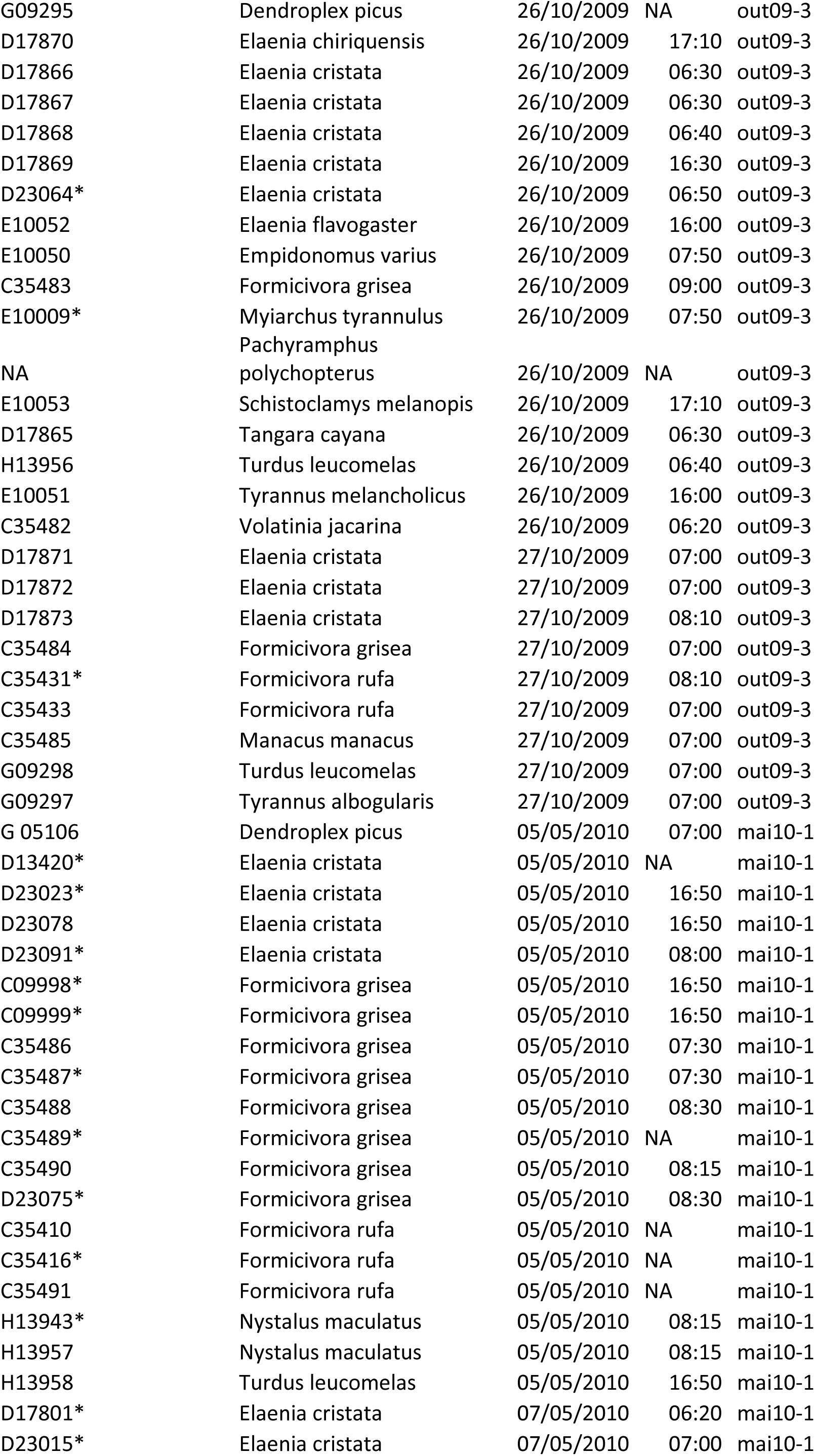
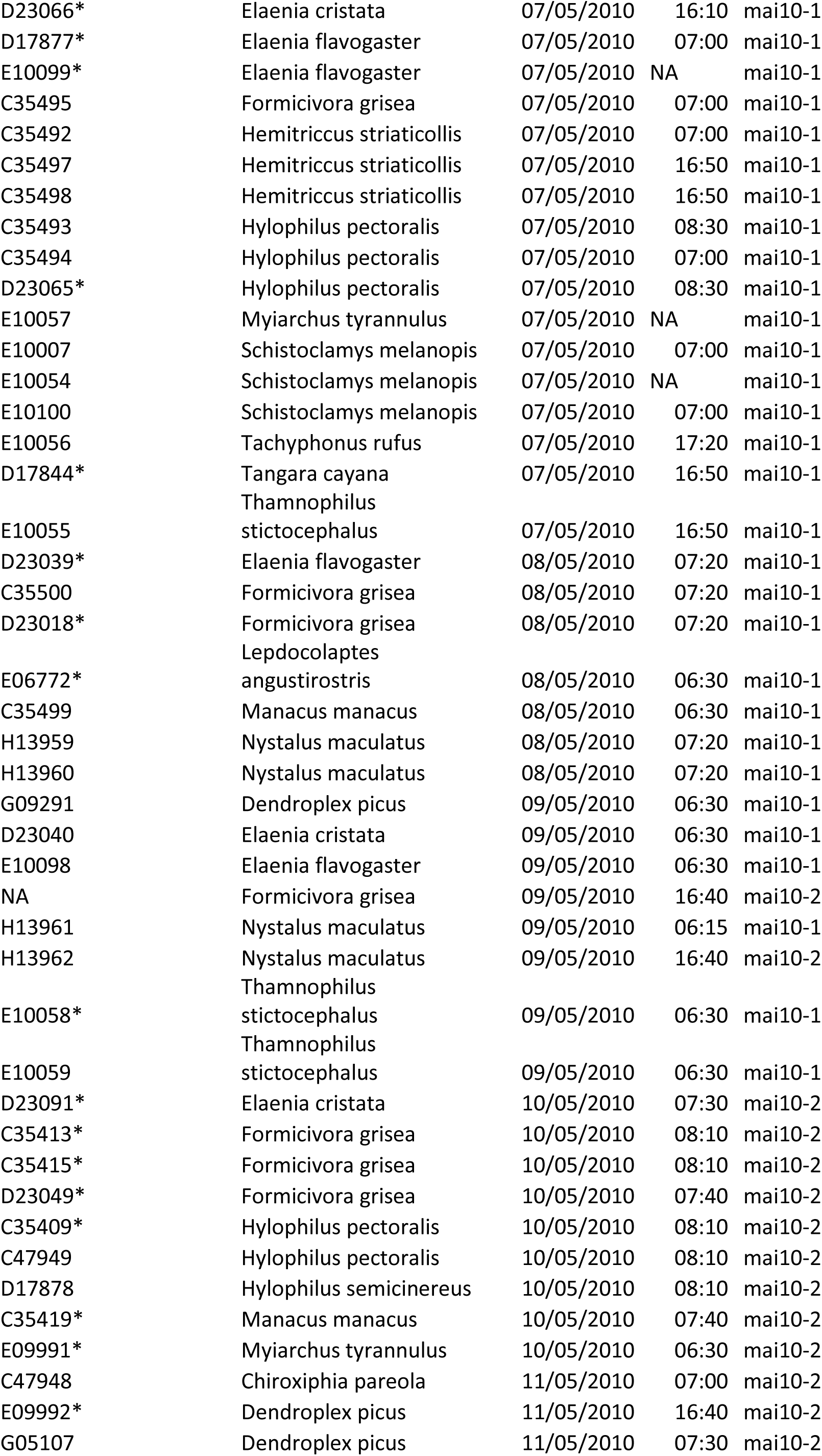
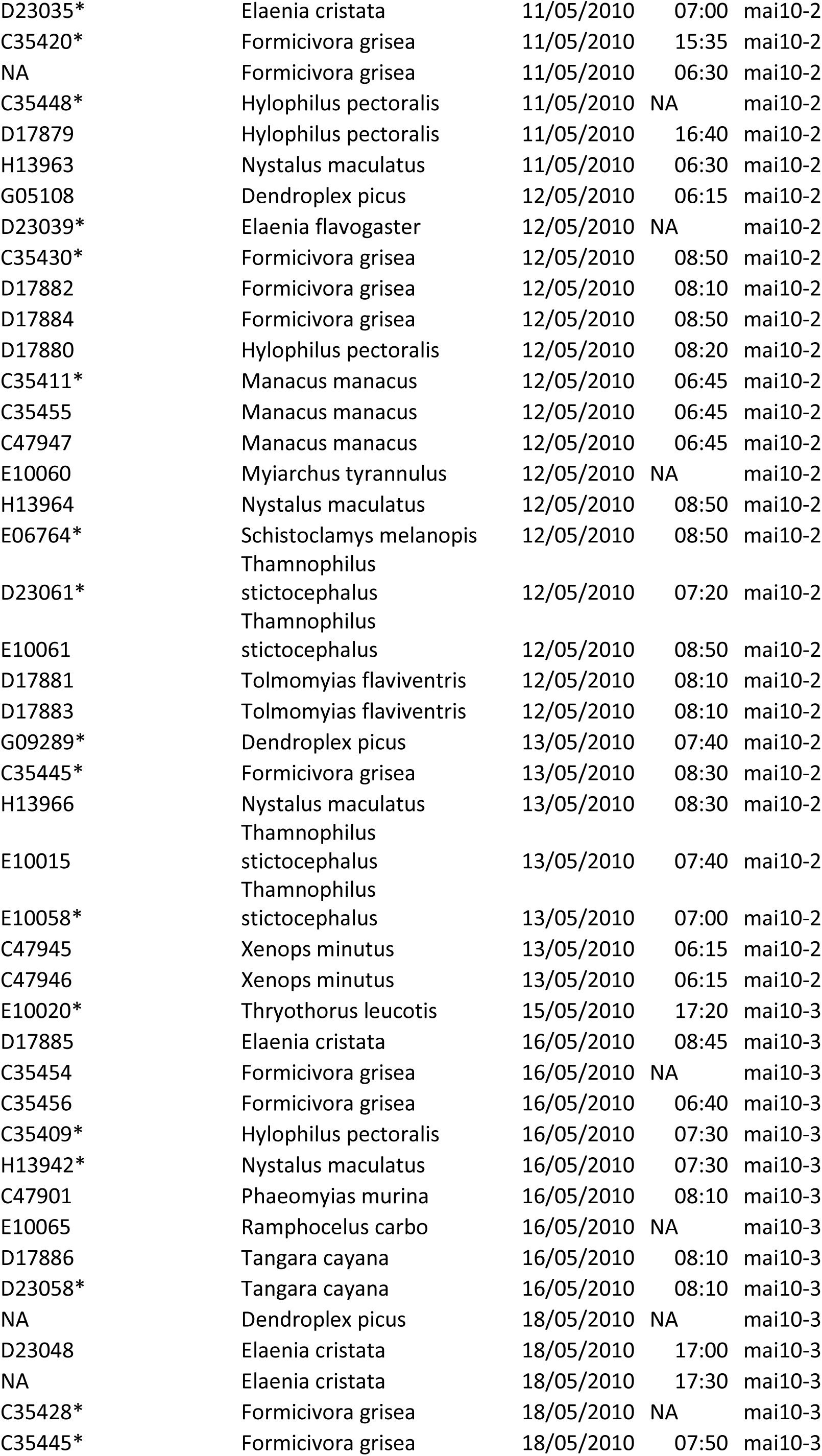
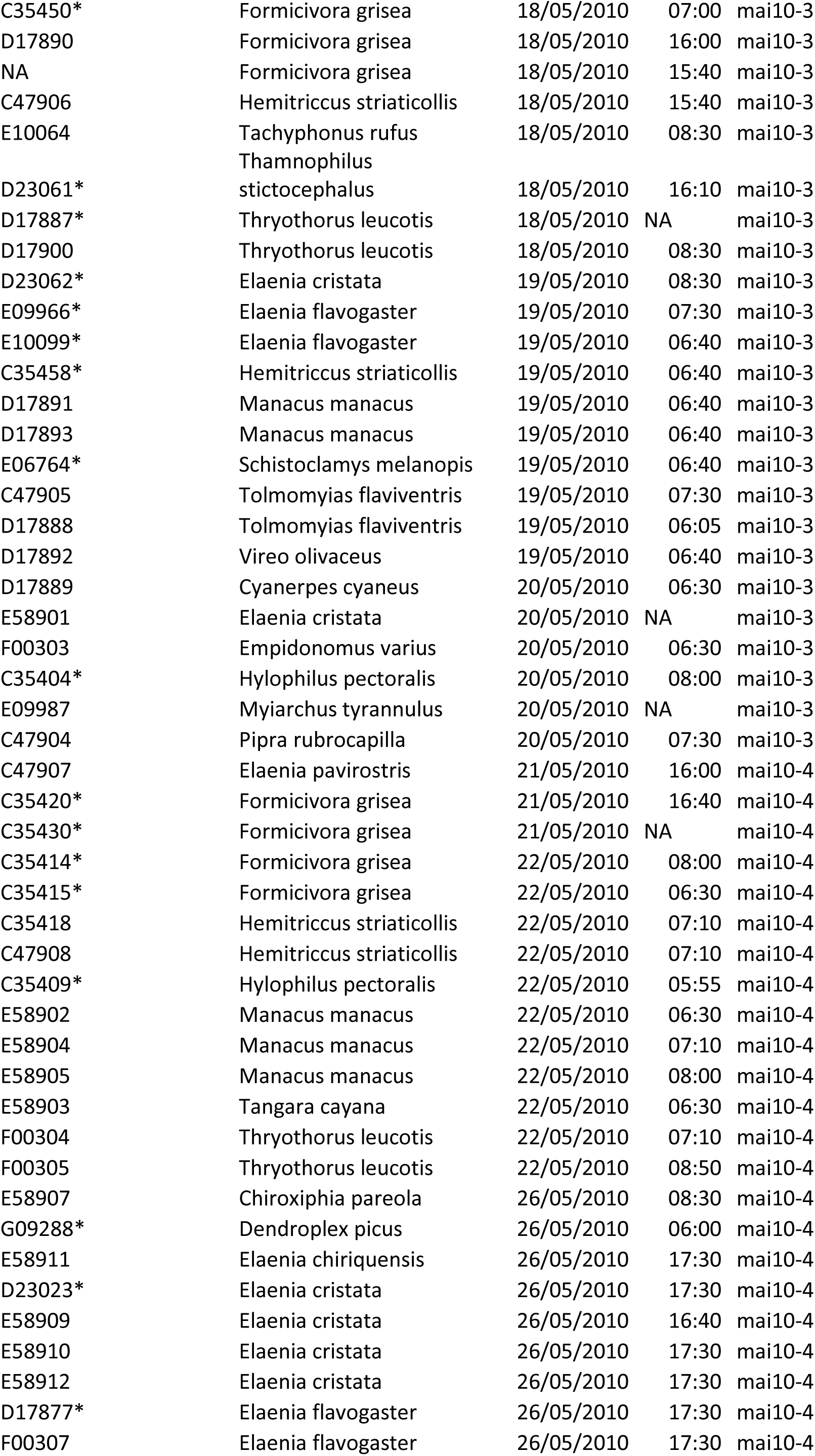
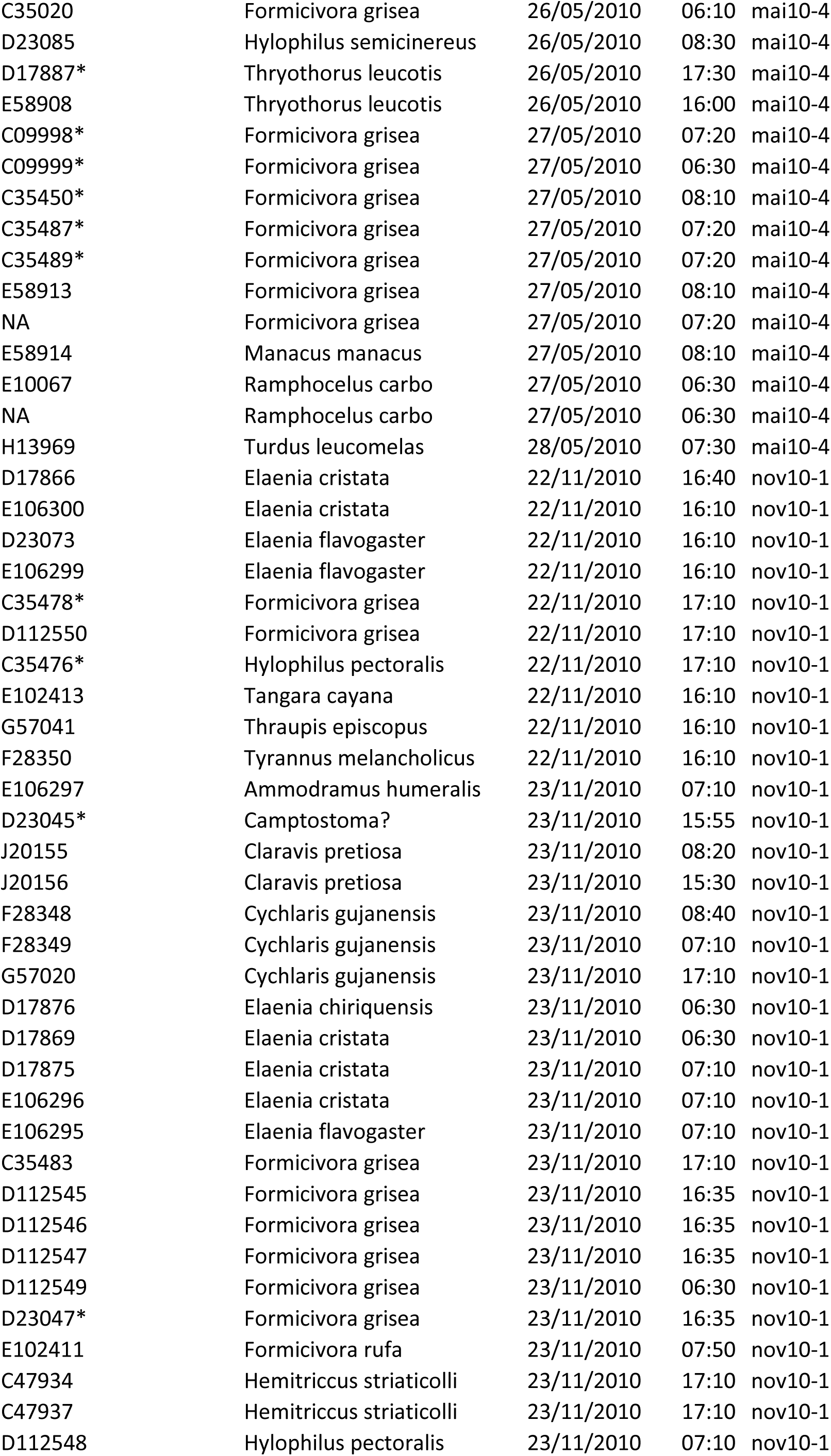
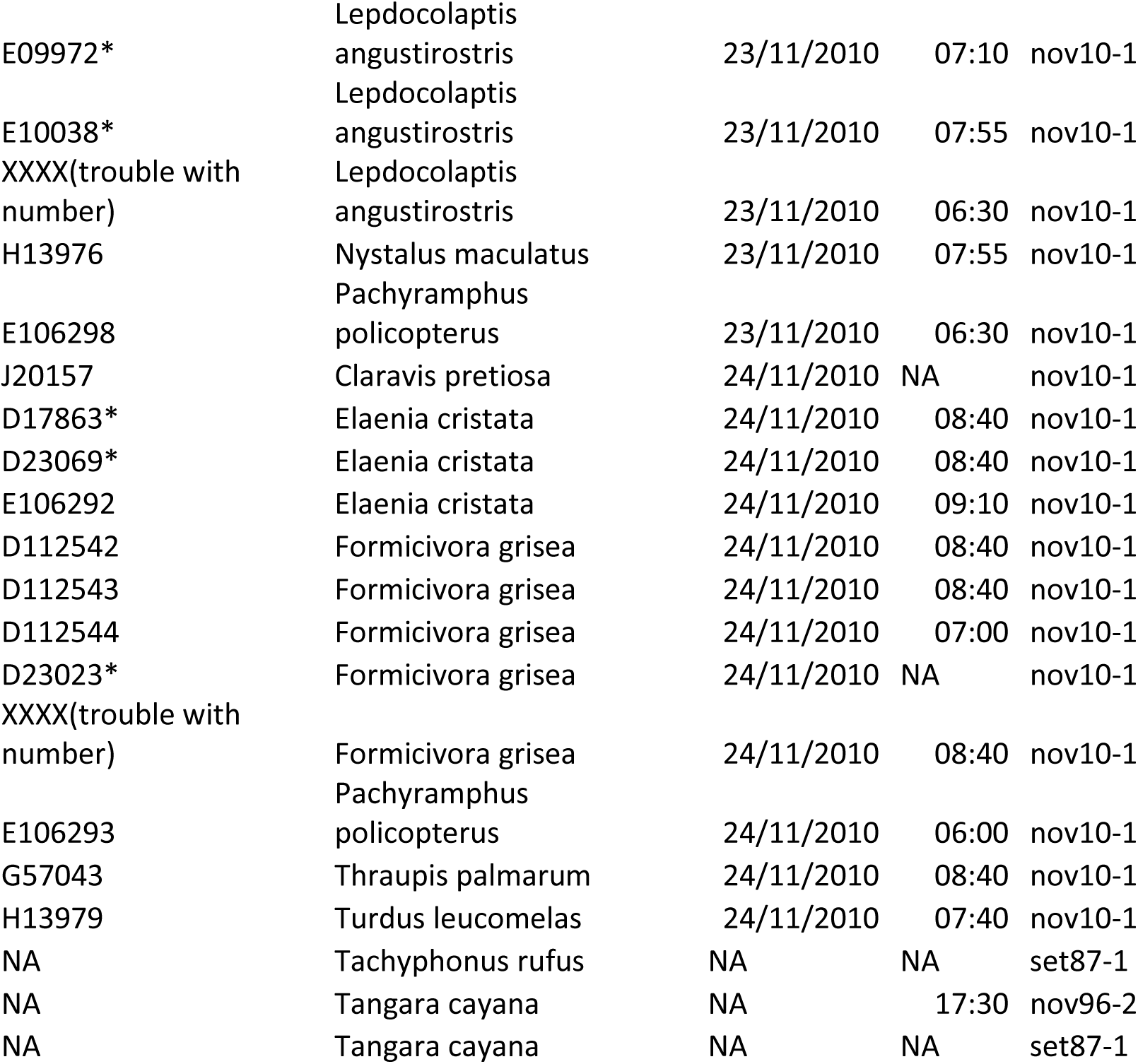
Birds captured in the temporal-analysis study. Birds with recaptures are marked with an “*”.

**Table S2:** Correlations among environmental variables. The upper diagonal shows the p-values and the lower diagonal the Pearson correlation.

|  | **Fire**<br>**frequency** | **Fire**<br>**extension** | **Tree**<br>**coverage** | **Forest**<br>**distance** |
| --- | --- | --- | --- | --- |
| **Fire**<br>**frequency** |  | <0.001 | 0.002 | 0.13 |
| **Fire**<br>**extension** | 0.96 |  | 0.002 | 0.22 |
| **Tree**<br>**coverage** | -0.8 | -0.79 |  | 0.05 |
| **Forest**<br>**distance** | 0.46 | 0.38 | -0.57 |  |

**Table S3:** Pairwise Wilcox-test results for forest-species abundance in the different sampling periods.

|  | **2009/10** | **1987/88** |
| --- | --- | --- |
| **1987/88** | <0.001 | - |
| **1996/97** | 0.001 | 0.3024 |

